# Loss of Ehmt2/G9a function in zebrafish is associated with global deficiency in H3K9 dimethylation, misregulated cell cycle dynamics, and embryonic developmental delay

**DOI:** 10.64898/2026.04.05.716391

**Authors:** Tara E. McDonnell, Francesca Meda, Steven J. Deimling, Vincent Tropepe

## Abstract

Ehmt2 is a key H3K9 methyltransferase that regulates genome silencing and structural integrity during animal development. In addition to this canonical function, Ehmt2 has also been implicated in neural tissues mediating both direct and indirect transcriptional activation, and exon splicing, to facilitate proper neural cell differentiation and survival. Several germline loss-of-function animal models have been developed showing both conserved and divergent phenotypes that range from embryonic lethality to behavioural deficits in adult, fertile animals. Here, we generated the first maternal-zygotic *ehmt2* loss of function mutant in zebrafish using CRISPR-Cas9 mutagenesis. An assessment of the pattern of H3K9 methylation in mutant embryos by ChIP-seq indicates that there are aberrant levels of this repressive mark, including reduction in discrete 5’ non-coding regions of genes, but with no significant change in the overall pattern distribution of these marks across the genome. Global transcriptome and morphological analyses demonstrated that mutant embryos displayed greater variation in the timing of developmental progression that is, on average, slower compared to controls. Despite this, mutant embryos ultimately survive and are fertile. Through examination of progenitor cell dynamics and gene expression profiles, we found that the delay in embryonic development was associated with longer rates of S-M phases of the progenitor cell cycle in mutants leading to deficits in tissue growth. Finally, our data suggest a robust network of epigenetic regulators can potentially compensate for Ehmt2 loss of function and permit embryonic development and survival in *ehmt2* mutant zebrafish. Our work establishes a zebrafish *ehmt2* loss of function model that will facilitate examination of the complex and varied roles of Ehmt2 in vertebrate development.

## INTRODUCTION

Regulation of cellular differentiation and tissue growth in early development is driven by an interplay of epigenetic modifications which influence gene expression, DNA replication and repair, and genome structure [1]. A key protein complex primarily responsible for catalysing early repressive modifications involves Euchromatic histone lysine methyltransferase (Ehmt2) (also known as G9a), which is known for depositing a dimethylation mark at Histone 3 Lysine 9 (H3K9me2) [2,3]. H3K9me2 is predominantly deposited in intergenic regions, including at repetitive sequences and transposable elements, and is associated with silenced regions in euchromatin [4]. Functionally, H3K9me2 regulates gene expression in several ways depending upon its localisation. While, H3K9me2 localisation at gene promoters/enhancers demonstrates a more straightforward repressive mechanism, H3K9me2 enrichment at transposable and repetitive elements can promote changes to the local environment, preventing expression of specific gene variants [5]. In addition to specific gene repression, global H3K9me2 has been shown to be critical for genomic integrity[6], and can be employed by cells as a protective mechanism in replication stress-induced DNA Damage [7].

Research over the last decade has indicated that Ehmt2 possesses a multifaceted role across specific tissues, that extends beyond its canonical role in gene silencing, including transcriptional activation and exon splicing. For instance, Ehmt2 is required in neural tissue to encourage proper cell differentiation and survival [8–11]. It has also been implicated in DNA replication and repair processes. In proliferating cells, Ehmt2 can deposit H3K56me1 which is recognised by proliferating cell nuclear antigen (PCNA) to promote DNA replication [12]. In response to DNA damage, Ehmt2 has been shown to recruit repair machinery, specifically 53BP, BRCA and RPA [13,14].

Current loss of function models of Ehmt2 have varying degrees of severity in organism development. Mouse germline genetic loss of function models for Ehmt2 (and its partner protein, Ehmt1) display a global loss of H3K9me2, and are embryonic lethal between embryonic days 9.5-12 due to severe growth defects [2,15]. In contrast, *Drosophila*, which possess a sole homologue of Ehmt1/2 (referred to as EHMT), are adult viable and fertile [16]. Unlike in mice, *Drosophila* mutants do not exhibit global loss of H3K9me2, but instead show a reduction of H3K9me2 specifically at neuronal and behavioural genes which correlates with behavioural abnormalities [17,18]. *C. Elegans* possess two known H3K9 methyltransferases: SET-25 (Ehmt2 homologue) that primarily catalyses H3K9me3, and MET-2 (SETDB1 homologue) that is required for H3K9 mono- and di-methylation [19]. SET-25 deficient worms show no viability/sterility defects, while MET-2 loss of function results in progressive, transgenerational loss of fertility that eventually leads to sterility. Furthermore, MET-2;SET-25 double mutants are able to progress through development but are adult sterile [19].

Despite significant research into Ehmt2 across various species and tissues, many questions remain, particularly surrounding the conserved or divergent functions of Ehmt2 in embryonic survival and development, and in what contexts Ehmt2 may be dispensable for growth and survival. Here, we investigate how *Ehmt2* germline loss of function affects early zebrafish development using a novel maternal-zygotic mutant model. We found reduced H3K9me2 levels across the genome, especially in discrete 5’ non-coding regions in genes, in this model using chromatin immunoprecipitation sequencing (ChIP-Seq) and immunofluorescent staining. A comprehensive analysis by RNA sequencing (RNA-Seq) indicated altered cell cycling and delayed development. Microscopy and flow cytometry corroborated these findings, demonstrating global developmental delay and tissue growth defects at embryonic stages, which is consistent with mouse models. Nonetheless, like the *Drosophila EHMT* mutants and *C. elegans SET-25* mutants, the zebrafish e*hmt2* maternal-zygotic mutants are adult viable and fertile.

## RESULTS

### *Ehmt2* mutant zebrafish are adult viable and fertile despite reduction in H3K9me2 expression

To examine the function of Ehmt2 in early embryonic zebrafish development, we established an *ehmt2* mutant model using a CRISPR-Cas9 gene targeting approach. Wildtype (WT) embryos were injected into the cell at the 1-cell stage with guide RNA targeting exon 5 of the *ehmt2* gene. Founders with mutations in the target region were outcrossed with WT embryos, and the resultant embryos raised to adulthood and in-crossed to produce two stable lines carrying specific mutant alleles. Here, we focus on one mutant line which possess a 4bp deletion in exon 5 which is predicted to result in a premature termination codon (herein denoted as *ehmt2^Δ⁴^*). This mutation is outlined in Figure 1A. We were able to raise homozygous mutants to adulthood and in-cross them to produce viable embryos. Furthermore, we observed no obvious post-embryonic phenotype (Figure 1B).

**Figure 1:**
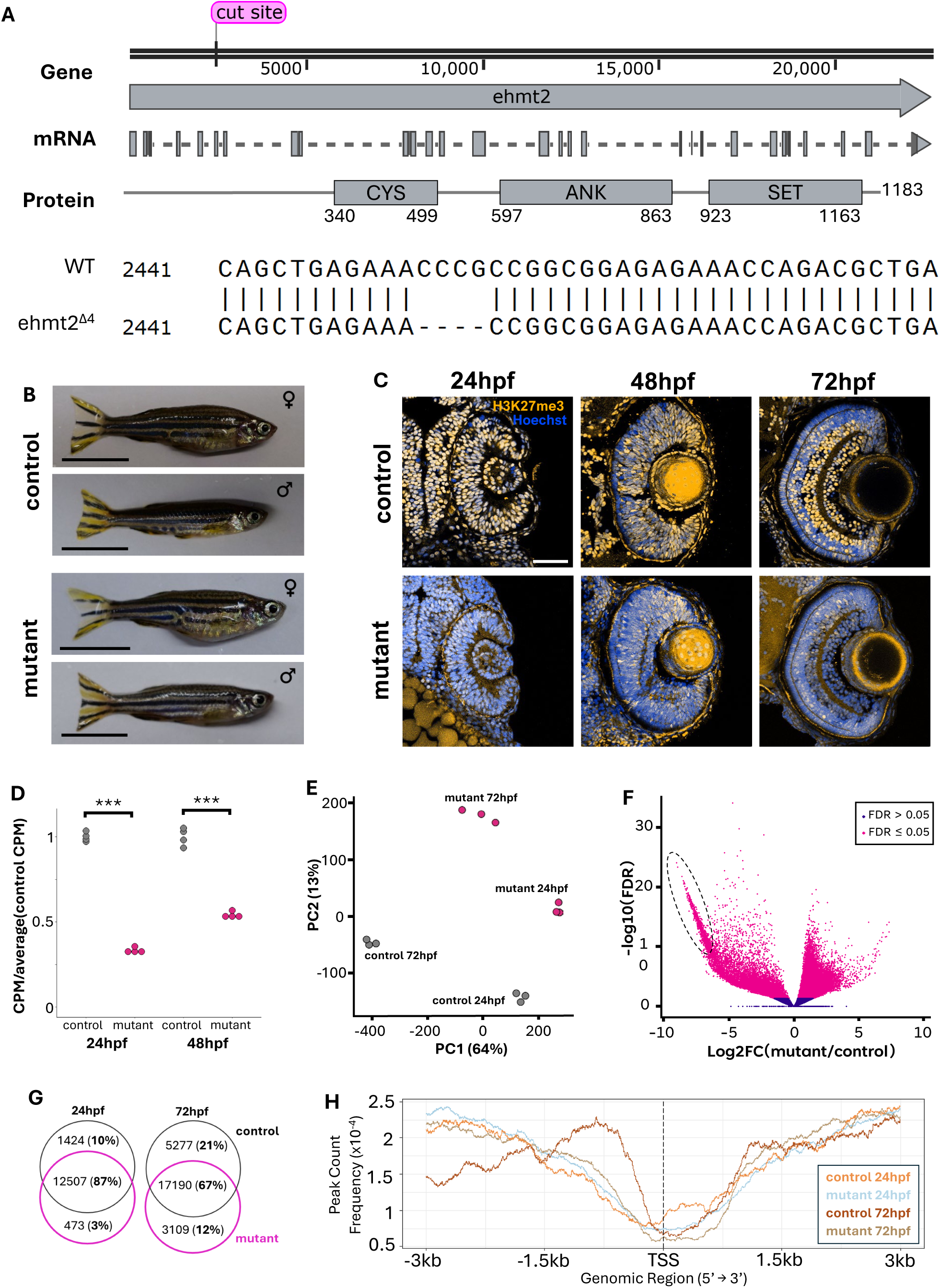
*Ehmt2* mutant embryos exhibit reduced H3K9me2 expression. (A) Zebrafish *ehmt2* gene locus, mRNA and protein structures, and DNA sequence showing the location of the 4bp deletion in *ehmt2^Δ⁴^* fish generated *via* CRISPR-Cas9. (B) Adult (15 mpf) control (top) and mutant (bottom) zebrafish. (C) Immunohistochemistry against H3K9me2 in retinal tissue (transverse sections) showed tissue-specific H3K9me2 loss at the cellular level. Scale bar = 50 µm, blue = Hoechst, yellow = H3K9me2. (D) *ehmt2* expression in whole embryos (RNA-Seq) at 24 and 48hpf plotted as CPM over the averaged CPM of control samples at that timepoint (n=4, *** = P ≤ 0.001). (E-H) Whole embryo ChIP-Seq against H3K9me2 (n = 3 biological replicates per group). (E) Principal component analysis confirms distinct clustering of mutant and control gene expression profiles at 24 and 72hpf. (F) Volcano plot of differentially enriched peaks in pink (|log2FC| ≥ 1 and P-value ≤ 0.05). (G) Venn diagrams showing overlap of differentially enriched peaks (|log2FC| ≥ 1 and P-value ≤ 0.05) between mutant and control embryos at 24hpf (left) and 72hpf (right). Percentage of differentially enriched peaks within the timepoint are in brackets. (H) Line graph indicating peak enrichment around transcription start sites across the genome. *Ehmt2* mutants had reduced H3K9me2 peaks within putative promoter regions at 72hpf (between 0-1500bp upstream of TSS) compared to control. FC = fold change, FDR = false discovery rate.

To confirm *ehmt2* loss of function, we examined *ehmt2* mRNA expression levels by analysing RNA-Seq data that were collected on 24 and 48 hours post fertilisation (hpf) pooled whole embryos. Figure 1D shows a significant reduction in *ehmt2* transcript levels in *ehmt2^Δ⁴/Δ⁴^* embryos compared to control at both timepoints (∼67% at 24hpf and ∼46% at 48hpf), suggesting that the mutant transcripts may be partially undergoing nonsense mediated decay. We also performed immunofluorescent analyses on sectioned retinal tissue between 24-72hpf and observed a global reduction of H3K9me2 at all three timepoints (Figure 1C). Moreover, similar reductions in H3K9me2 staining were observed in wildtype embryos treated with specific Ehmt2 pharmacological inhibitors (Figure S1A) or in wildtype embryos injected with an *ehmt2* gene specific morpholino (Figure S1B), consistent with our mutant loss of function data.

To further examine the effect of *ehmt2* loss of function on the distribution of H3K9me2 marks across the genome in our mutant, we performed ChIP-Seq against H3K9me2 on pooled whole *ehmt2^Δ⁴/Δ⁴^* and controls for 24 and 72hpf embryos. Principal component analysis (PCA) showed distinct clusters for each genotype and timepoint (Figure 1E). The volcano plot in Figure 1F highlights a population of sites with significant reduction in H3K9me2 in *ehmt2^Δ⁴/Δ⁴^* embryos (circled data points). When we interrogated H3K9me2 enrichment sites across the genome, we found that the majority of sites were located in distal, intergenic regions, consistent with previously published data, and this overarching distribution was not altered in the mutant compared to control (Figure S2) [5]. However, for peaks surrounding gene loci, we found that control embryos showed a shift in enrichment upstream of gene start sites (0-1500bp) at 72hpf compared to 24hpf, but this developmental shift in enrichment is absent at 72hpf in *ehmt2^Δ⁴/Δ⁴^* embryos (Figure 1H). When we compared the peak site overlap in *ehmt2^Δ⁴/Δ⁴^* and WT control embryos, we found less overlap at 72hpf compared to 24hpf (Figure 1G). These findings indicate that *ehmt2^Δ⁴/Δ⁴^* embryos can establish H3K9me2 marks at some early timepoint (prior to 24hpf), but they cannot efficiently deposit H3K9me2 at least at a portion of new sites as embryonic development proceeds to later stages (72hpf).

### The transcriptome profile of *ehmt2* mutants suggests impaired developmental timing

To examine how gene expression was affected in Ehmt2 loss of function mutants, we analysed our RNA-Seq data on 24 and 48hpf mutant and WT embryos. Clustering from PCA indicated that there were global differences in expression profiles between mutant and WT control embryos and that these differences were greater at the latter timepoint (Figure 2A), which aligns with the findings of our ChIP-Seq data. Figure 2B depicts a volcano plot highlighting the subset of genes differentially expressed in *ehmt2^Δ⁴/^*^Δ⁴^ compared to control. Differentially expressed genes were hierarchically clustered using Clust [20] which produced two distinct gene expression patterns (Figure 2C). Gene Ontology (GO) analysis was then employed on the clustered genes to identify differentially enriched pathways (Figure 2D). The analysis showed that genes involved in neural, regulatory, and developmental pathways remained upregulated in *ehmt2^Δ⁴/Δ⁴^* at 48hpf, when they would normally be downregulated in control embryos. The other expression pattern showed genes involved in cell cycling, cell metabolism and DNA replication & repair were downregulated in *ehmt2^Δ⁴/Δ⁴^* embryos at 24hpf and were further downregulated compared to control embryos at 48hpf.

**Figure 2:**
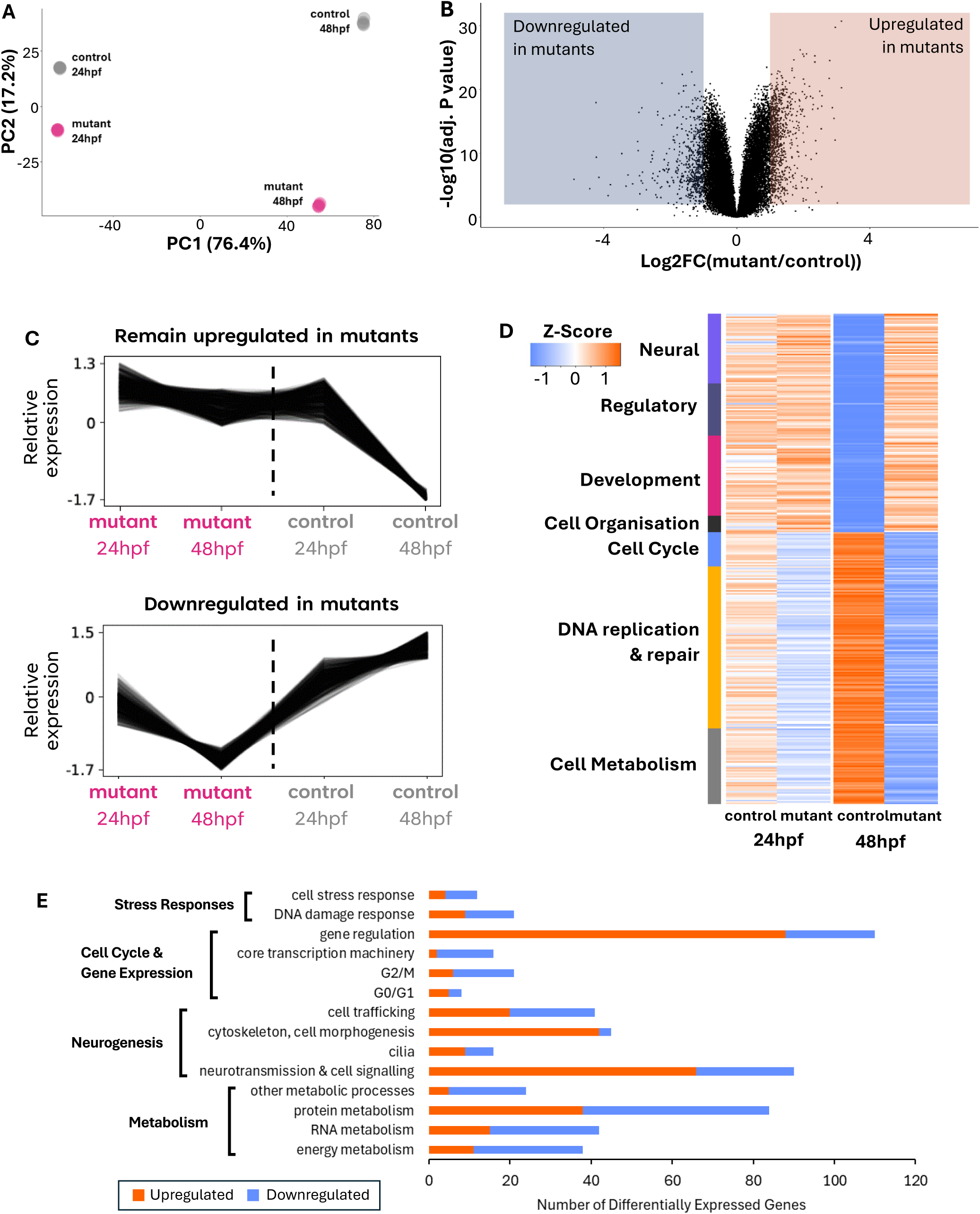
Differential gene expression in *ehmt2* mutants compared to control through whole-embryo RNA-Seq. (A) Principal component analysis confirms distinct clustering of mutant and WT control samples at both 24 and 48hpf. (B) Volcano plot highlighting differentially expressed genes (|log2FC| ≥ 1 and P-value ≤ 0.05). (C) Unsupervised hierarchal clustering of differentially expressed genes produced two distinct clusters of expression patterns. (D) Heatmap of differentially expressed genes represented in (C), sorted using Gene Ontology (GO) enrichment analysis. Genes involved in neural, regulatory and developmental processes remained upregulated in mutants at 48hpf compared to control, while genes involved in cell cycling, cell metabolism and DNA damage response were downregulated at both 24 and 48hpf in *ehmt2* mutants compared to control. (E) Plot indicating the number of identified DEGs sorted into functional subcategories. Orange = genes upregulated in mutants, blue = genes downregulated in mutants. n = 4 replicates per group.

By analysing subcategories, we found that a large number of genes involved in DNA replication and repair were dysregulated (Figure 2E) in the mutants, consistent with aberrant H3K9me2 deposition [13,21–24]. The downregulation of these genes and other cell cycle/metabolism pathways, along with subtle differences observed in 24-72hpf mutant embryonic growth, led us to speculate that global processes of development and/or growth are affected in *ehmt2^Δ⁴/Δ⁴^* embryos. Furthermore, of the upregulated genes identified at 48hpf relative to controls, 176 out of 350 (∼50%) are known to function in neural pathways including neurogenesis and neurotransmission, indicating that neurodevelopment genes that are normally downregulated between 24hpf and 48hpf in WT embryos are abnormally sustained in the mutants at 48hpf. We hypothesized that the observed changes in global developmental gene expression and H3K9me2 distribution in the *ehmt2^Δ⁴/Δ⁴^* embryos may, in part, be a result of developmental delay and/growth deficits in the mutant embryos.

### *Ehmt2* loss of function results in developmental delay

To examine how ehmt2 loss of function affects early developmental timing in zebrafish, we staged *ehmt2^Δ⁴/Δ⁴^* and WT control embryos at the 4-cell stage and measured how long these embryos took to reach Prim-5 (typically reached at ∼24hpf). Our data indicate that *ehmt2^Δ⁴/Δ⁴^* mutants have a delay in embryonic development, appearing by the Sphere stage, and persisting through to prim-5 (Figure 3A). By prim-5, the mutants are, on average, 3 hours behind WT control embryos. Our results also showed increased intra-clutch timing variation in mutants at all stages measured except for the earliest timepoint tested (128-cell stage). The variation in developmental timing to Sphere stage is shown in Figure 3B, with all other developmental stages shown in Figure S3. These data suggest that Ehmt2 may be involved in suppressing variable developmental delay in zebrafish, which is consistent with previously published data [4].

**Figure 3:**
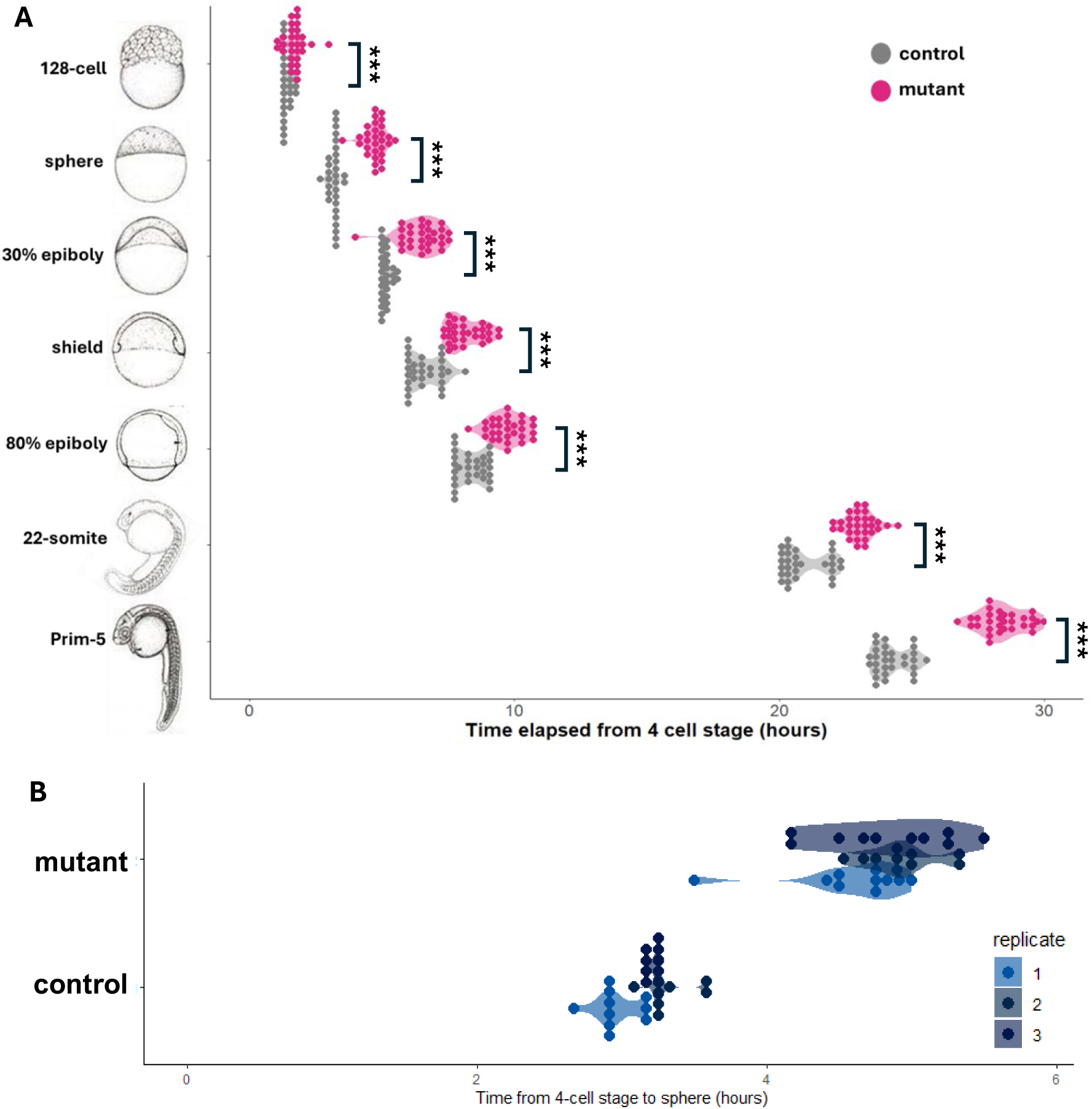
*Ehmt2* mutant embryos display an early developmental delay. (A) Plot depicting time elapsed since the 4-cell stage for mutant (pink) and control (grey) embryos to reach certain developmental stages (n = 30 per group). Each stage tested showed significant (P ≤ 0.001) variation between mutant and WT embryos. B) Plot of time elapsed for embryos to reach sphere stage from the 4-cell stage (from A), coloured by replicate, showing increased intra-clutch variation.

Since our RNA-Seq analysis indicated that Ehmt2 loss of function had a negative impact on the normal progression of neural development, and based on the previously reported roles of Ehmt2 in nervous system development and function in mice [8,10,25,26], we next wanted to examine whether we could detect abnormal growth of neural tissue in our mutants, focusing on the development of the embryonic retina. We stage-matched embryos at prim-5, then whole-mounted and stained with Hoechst (nuclear stain) to compare retinal size in mutant and control embryos. Representative images are shown in Figure 4A, and retinal size is quantified in Figure 4B-C. We found that *ehmt2^Δ⁴/Δ⁴^* retinas were significantly smaller than stage matched WT control retinas at prim-5, however, when we measured retinal size two days after prim-5 (normally around 72hpf in WT embryos), the size difference was negligible. These data show that *ehmt2^Δ⁴/Δ⁴^* embryos possess an early developmental delay, and that retinal tissue growth is significantly reduced at prim-5 (compared to control) but seems to reach a comparable size by 72hpf.

**Figure 4:**
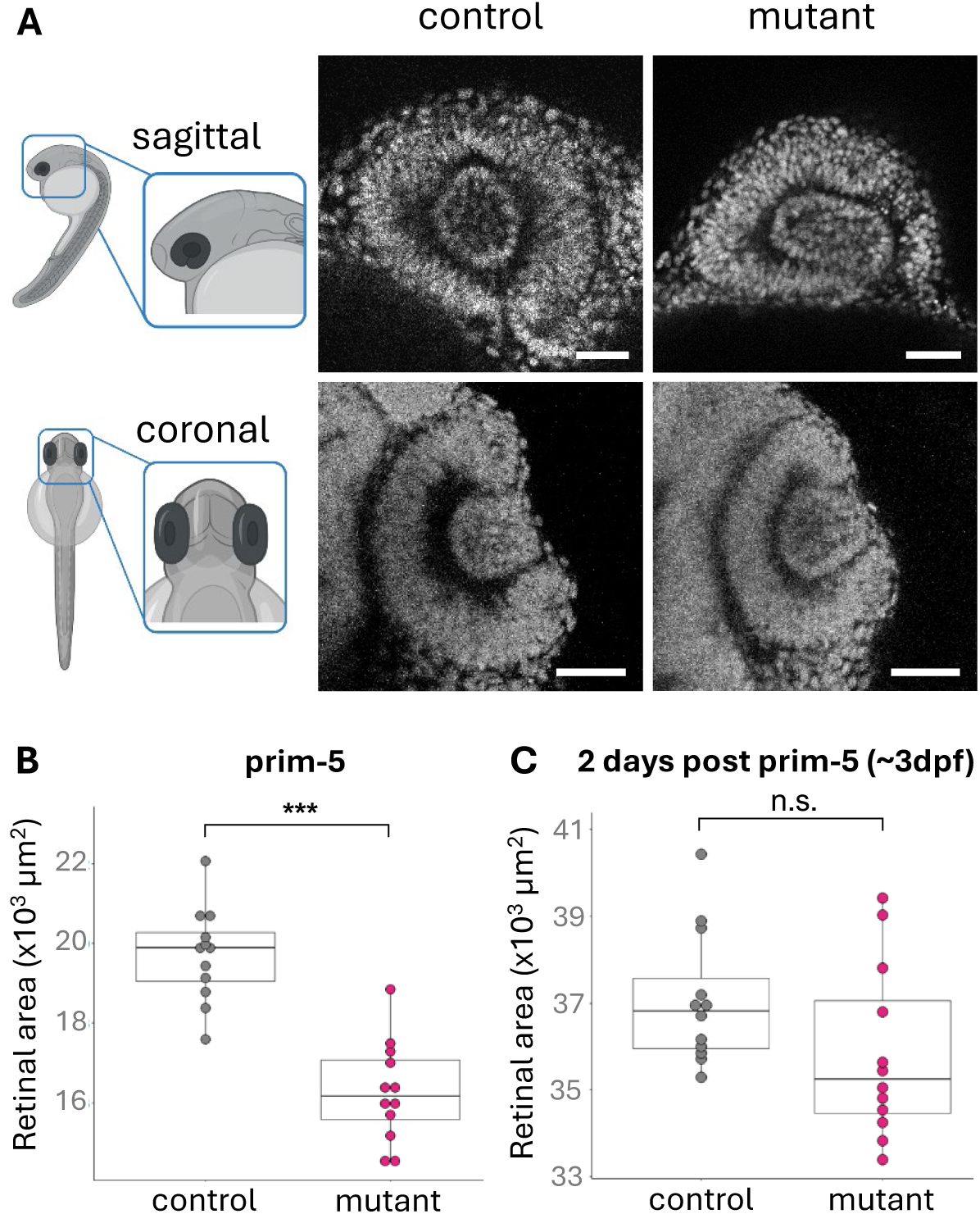
*Ehmt2* mutant retinas are smaller at prim-5, but the reach comparable sizes by around 3dpf. (A) Representative whole-mount images of control (left) and mutant (right) retinas stage matched at prim-5, stained with Hoechst. Scale bars = 50 μm. (B) Retinal cross-sectional area at prim-5 showing significant size difference between mutant and control embryos. (C) Retinal cross-sectional area measured two days post prim-5 showed the observed size difference is no longer significant between mutant and control embryos. Retinal cross-sectional areas were measured from coronal z-stacks through the central region of the retina. *** = P ≤ 0.001, n.s. = not significant. n = 12 per group.

### Ehmt2 is required for normal cell cycle timing during early zebrafish development

Our findings indicated that *ehmt2^Δ⁴/Δ⁴^* embryos were developmentally delayed at both a whole-embryo, and tissue-specific level. To address how Ehmt2 loss of function results in this delay, we first assessed whether the mutant tissues displayed abnormally higher levels of cell death. Focusing on the retina, we show using Acridine Orange [27] in live mutant and WT embryos there were no differences in cell viability at either 24hpf or 48hpf (Figure S4). These findings were validated by staining retinal sections using an antibody against cleaved caspase-3 (data not shown). Following this, we used immunofluorescent staining on retinal sections to detect proliferating cell nuclear antigen (PCNA) and phospho-histone H3 (PHH3) expression from 24-72hpf, to assess whether there were any changes in the proportion of cells engaged in the cell cycle (PCNA+) or specifically in mitosis (PHH3+) in the embryonic retina. We found that there was an increase in the proportion of PCNA expressing cells at both 24hpf and 48hpf in *ehmt2^Δ⁴/Δ⁴^* embryos, but not at 72hpf (Figure 5A, C). In contrast, the relative proportion of PHH3+ cells was significantly reduced at 24hpf, and greater at 48hpf, in the mutants, but again no differences were observed at 72hpf (Figure 5B, D).

**Figure 5:**
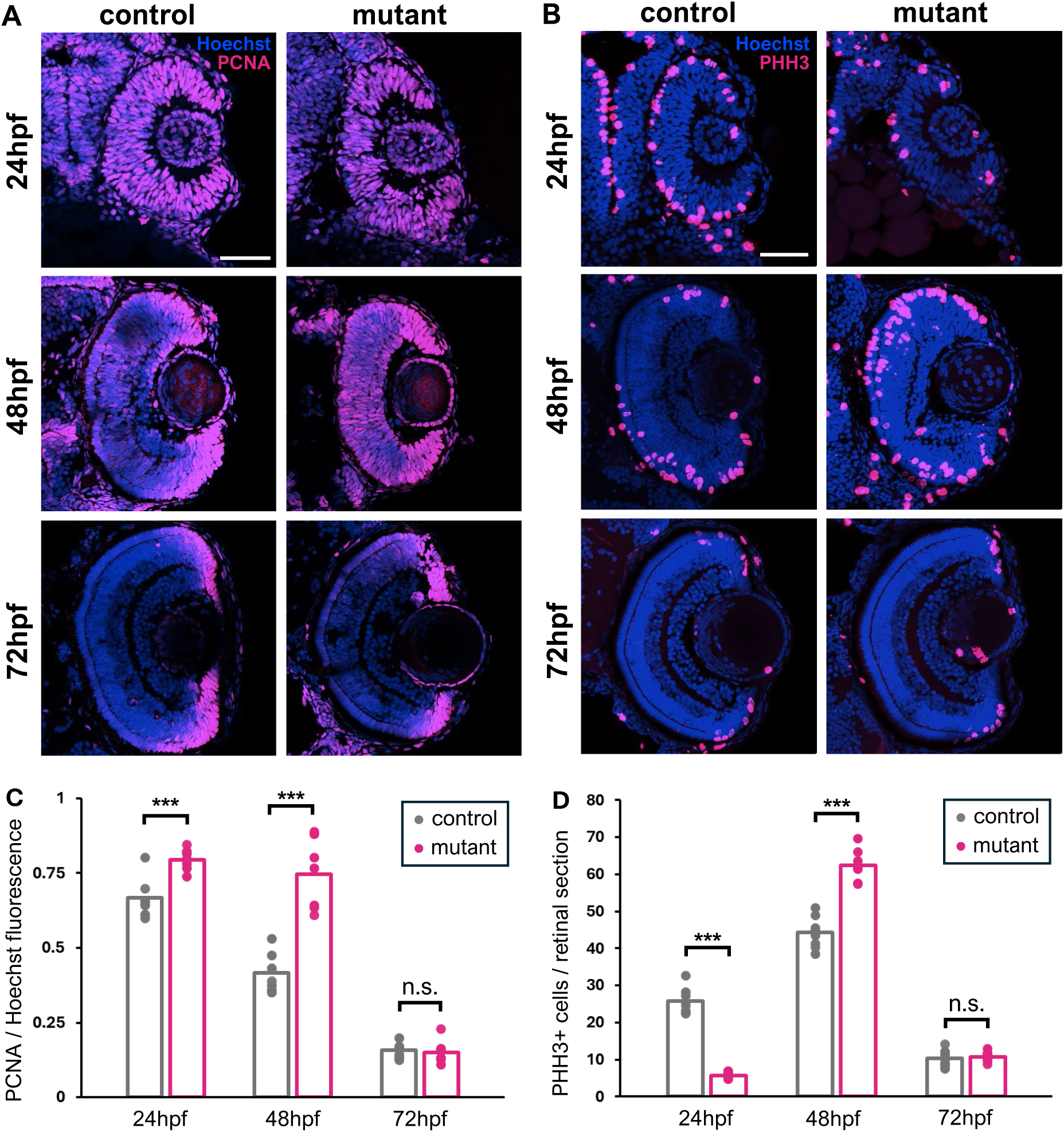
Retinal Proliferation persists beyond normal developmental timing. Proportion of proliferating cells is increased in mutant retinas at early timepoints but matches control by 72hpf. Immunostaining against PCNA (A) and Phospho-Histone H3 (B) in retinal sections with quantification of fluorescence shown in (C) and (D), respectively. Bar plots indicate average per sample type. Scale bars = 50 µm, n = 8 per group, *** = P ≤ 0.001, n.s. = not significant, error bars = 1 standard deviation.

It is possible that the apparent dynamic changes in the proportion of cycling cells in the mutant are due to alterations in the duration of the different phases of the cell cycle in proliferating retinal progenitor cells. To examine this, we performed Flow Cytometry on cells obtained from 24hpf mutant and control (whole) embryos stained with DAPI (nuclear stain) to examine the proportion of cells in each cell cycle phase. Our results, shown in Figure 6A, revealed a small but significant increase in the proportion of cells in S-M phases of the cell cycle in mutants compared to WT control. These findings indicate that cells from *ehmt2^Δ⁴/Δ⁴^* embryos may spend slightly longer in these phases compared to control.

**Figure 6:**
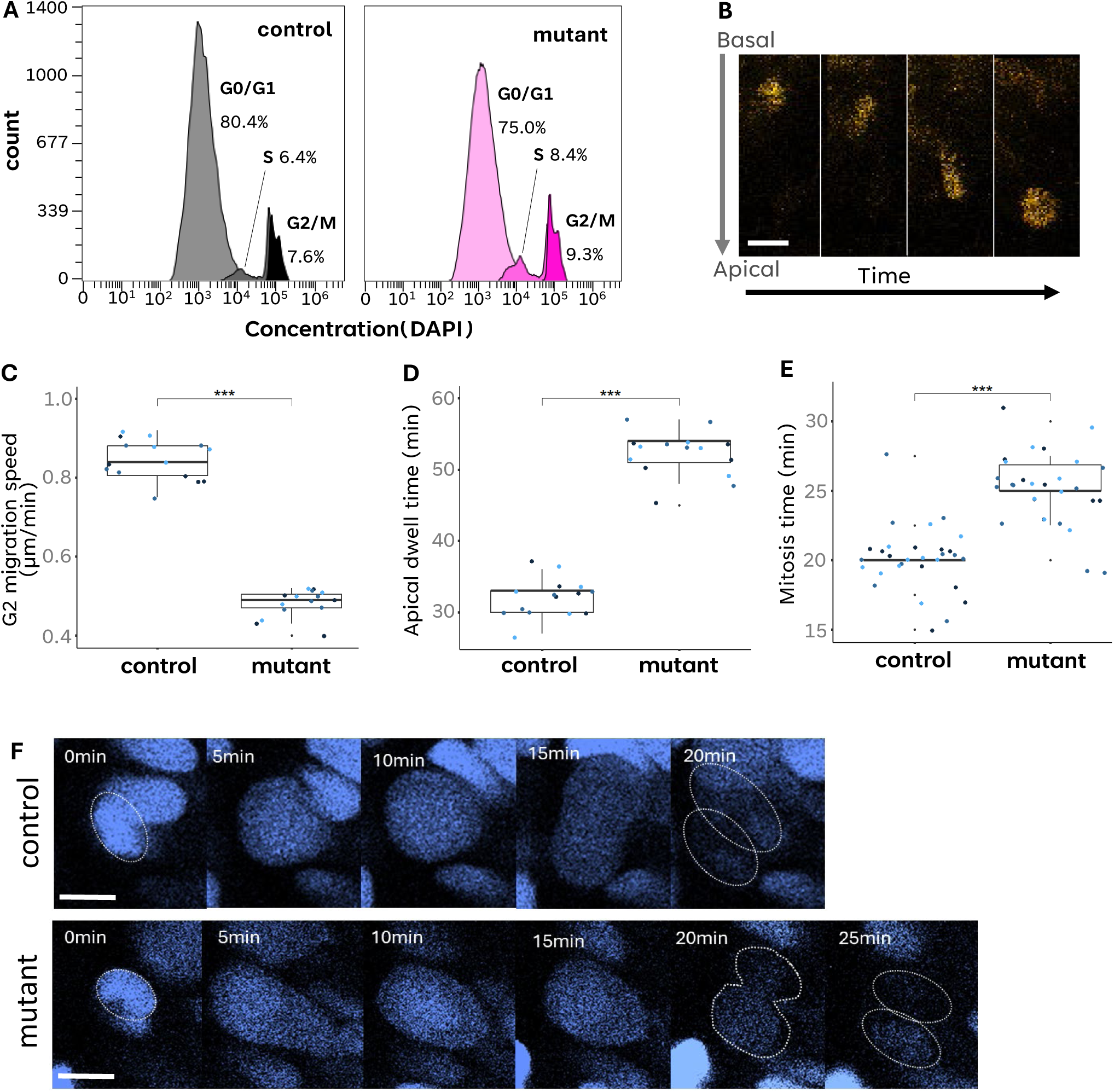
*Ehmt2* mutant embryos display prolonged S-M phases at 24hpf. (A) Cell cycle analysis *via* flow cytometry from pooled whole embryos at 24hpf (left = control, right = mutant). (B) A representative series of a single WT mAG-positive nuclei in G2 which undergoes interkinetic nuclear migration to the apical side of the developing retina. Imaged between 20-27 hpf, scale bar = 10 μm. (C) Migration speed of interkinetic nuclear migration during G2 in mutant and control retinal progenitor cells (n = 5 nuclei x 3 embryo replicates). (D) Dwell time of nuclei at the apical lamina before entering mitosis for mutant and control retinal progenitor cells (n = 5 nuclei x 3 embryo replicates). (E) A representative series of a single H2A:GFP-positive nuclei during mitosis in the mutant (lower) and control (upper) retina between 21-24 hpf. F) Mitosis time in mutant and control retinal progenitor cells (n = 9 nuclei x 3 embryo replicates, scale bar = 5 µm). *** = P ≤ 0.001.

To validate these findings directly, we turned to a live imaging approach using a transgenic line for *geminin* expression (*Tg[EF1a:mAG-zGem(1/100)^rw0410h]*), a gene upregulated during S phase, maintained in G2, and rapidly degraded after mitosis [28]. This transgenic line, hereafter reported as *Tg[zGem]*, was crossed to the *ehmt2^Δ⁴/Δ⁴^* mutants to obtain mAG-positive homozygous mutants and WT siblings. Embryos derived from these transgene-expressing *ehmt2^Δ⁴/Δ⁴^* mutants were agarose-mounted and imaged live using a confocal microscope from 20-27hpf allowing for direct visualisation of retinal progenitor cell cycle dynamics. A sample sequence of images showing migration of a WT nucleus moving apically in concordance with G2, followed by mitosis, is shown in Figure 6B (also see Figure S5). The speed of interkinetic nuclear migration was measured during G2 and we found a significant reduction in speed in *ehmt2^Δ⁴/Δ⁴^* embryos (Figure 6C). Furthermore, we observed that mutant nuclei had an increase in dwell time at the apical membrane prior to undergoing mitosis (the onset of mitosis was identified by nuclear condensation). The dwell time for nuclei in an *ehmt2^Δ⁴/Δ⁴^* mutant and a WT sibling is depicted in Figure 6D. To measure mitosis timing, the *ehmt2^Δ⁴/Δ⁴^* mutant fish were crossed to a separate transgenic reporter line for histone H2A (*ehmt2^Δ⁴/Δ⁴^; Tg(H2A:GFP)*) to fluorescently label all live nuclei and was bred to homozygosity. These embryos were imaged between 21-24hpf to show that time to completion for mitosis was significantly increased in progenitor cells of the *ehmt2^Δ⁴/Δ⁴^* retina compared to their WT siblings (Figures 6E, F). Taken together, our live imaging results indicate prolonged G2/M phases in individual progenitor cells within *ehmt2^Δ⁴/Δ⁴^* retinas, correlating with the population level changes in cell cycle dynamics in the mutant retina.

### Putative compensation in epigenetic regulatory processes may promote ***ehmt2****^Δ^****^4/^****^Δ^****^4^* embryonic survival**

We have shown that, while *ehmt2^Δ⁴/Δ⁴^* embryos are developmentally delayed, which is correlated with altered cell cycle dynamics in progenitor cells, and have altered global gene expression patterns, they are ultimately able to progress beyond embryogenesis and survive to adulthood. We thus examined possible compensatory pathways that might allow embryonic survival in *ehmt2^Δ⁴/Δ⁴^* mutants. Given, the crosstalk between many epigenetic networks, we examined the differential expression of epigenetic modifiers, as depicted in Figure 7A. Of the differentially expressed epigenetic modifiers identified in our RNA-Seq analysis, we found the majority of chromatin remodellers (Figure 7A, left panel), and histone markers associated with promoting active transcription (Figure 7A, middle panel), remained upregulated in *ehmt2^Δ⁴/Δ⁴^* embryos at 48hpf.

**Figure 7:**
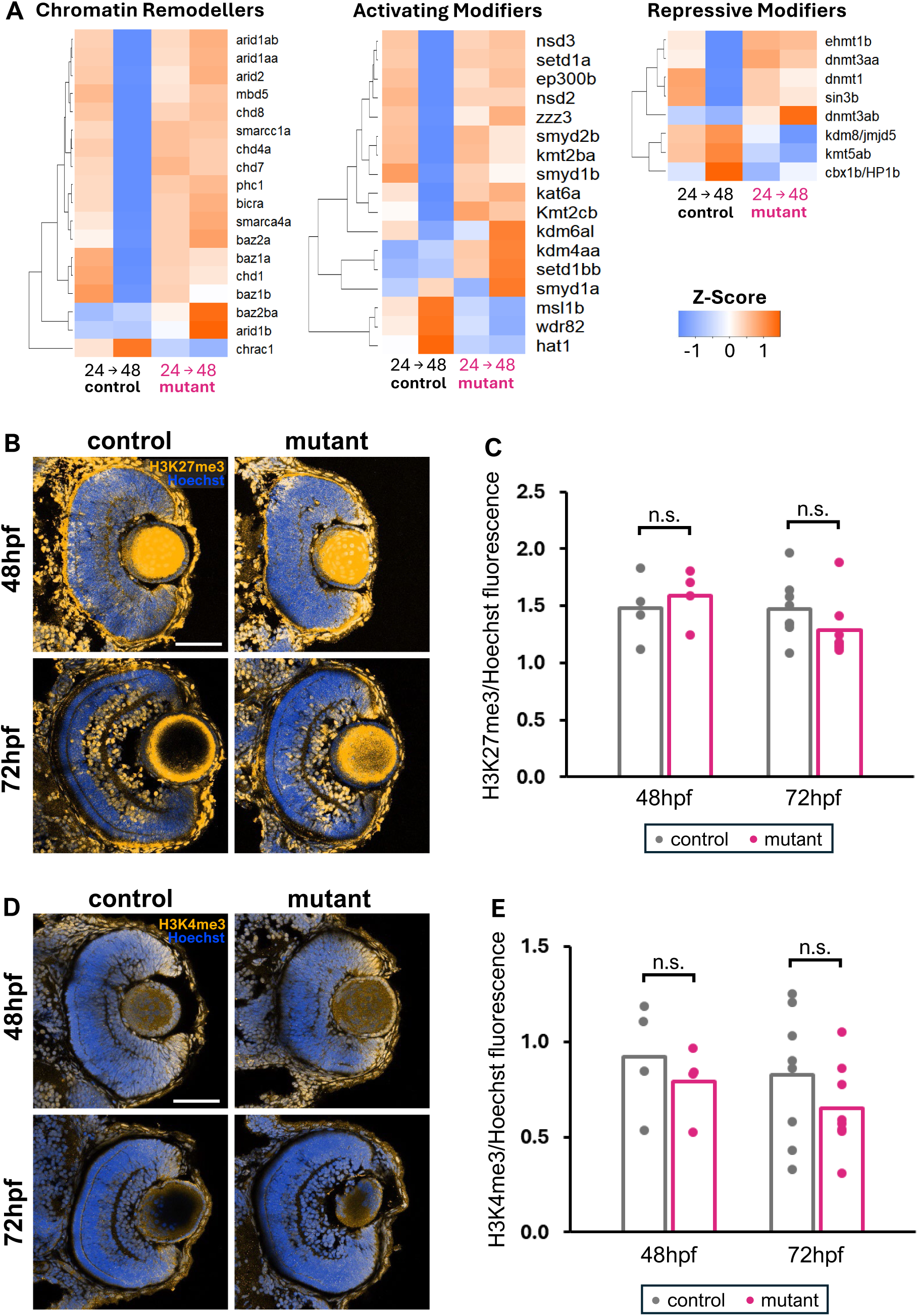
Expression of epigenetic modifiers and marks in mutants compared to controls. (A) Heatmaps showing expression patterns of DEGs with canonical roles in chromatin remodelling (left), activating epigenetic modifications (middle), and repressive epigenetic modifications (right) from whole embryo RNA-Seq. (B) Immunohistochemistry against H3K27me3 in retinal tissue (transverse sections). Yellow = H3K27me3, blue = Hoechst. (C) Fluorescence intensity quantification of H3K27me3 (from B) shows no significant difference in global H3K27 trimethylation enrichment between mutant and control retinas. (D) Immunohistochemistry against H3K4me3 in retinal tissue (transverse sections). Yellow = H3K4me3, blue = Hoechst. (E) Fluorescence intensity quantification of H3K4me3 (from D) shows no significant difference in global H3K27 trimethylation enrichment between mutant and control retinas. Bar plots indicate average per sample type. Scale bars = 50 µm; 48 hpf n = 4 per group; 72 hpf n = 8 per group; n.s. = not significant.

The expression of *ehmt1b*, an orthologue of Ehmt1 in mammals, which is a key co-regulator of Ehmt2 function [2,29], is upregulated at 48hpf in mutants relative to controls (Figure 7A, right panel), suggesting the possibility of transcriptional adaptation to compensate for the loss of Ehmt2 [30,31]. However, Ehmt1 has not yet been shown to be capable of dimethylating H3K9 *in vivo* [32], and our immunofluorescent data does not indicate that it is functionally compensating for global H3K9me2 deposition in the mutant. DNA methylation is another mark critical for gene repression. Interestingly, we found increased expression of DNA methyltransferases *dnmt1*, *dnmt3aa*, and *dnmt3ab* in *ehmt2^Δ⁴/Δ⁴^* embryos (Figure 7A, right panel), which could offer a potential compensatory mechanism for silencing in the absence of Ehmt2-mediated H3K9me2 deposition. However, given that Ehmt2 has been reported to be critical for Dnmt recruitment [33,34], further analysis is needed to determine whether increased Dnmt expression has a functional consequence in *ehmt2^Δ⁴/Δ⁴^* embryos.

Another key repressive mark is H3K27 methylation. Previously, work has shown distinct functions for H3K27 methylation and H3K9 methylation, where H3K27 methylation possesses a context-dependent, regulatory role, while H3K9 methylation is required for genome stability and DNA methylation [35]. While their functions and deposition sites do not always overlap, our RNA-Seq results did not show altered EZH1/2 (PRC1/2) expression (*data not shown*), which are required for H3K27me2/3. We also performed immunofluorescent staining for the presence of H3K27me3 marks in the embryonic retina and found no observable differences in global levels of H3K27me3 between *ehmt2^Δ⁴/Δ⁴^* and WT control retinas (Figure 7B-C). Taken together, these findings indicate H3K27me3 does not appear to be globally affected in *ehmt2^Δ⁴/Δ⁴^* embryos.

Finally, we examined a mark associated with active transcription, H3K4me3, since we found significant sustained upregulation of several genes associated with H3K4 methylation (e.g., *kmt2ba*, *kmt2cb*, *smyd2b* and *setd1a*) (Figure 7A, middle panel). We compared the levels of H3K4me3 marks in mutant and control retinas using immunofluorescence, but similar to the H3K27me3 results described above, there were no significant differences in global H3K4me3 (Figure 7D-E).

Combined, our results show that while we identified gene expression changes to various histone modifiers in *ehmt2^Δ⁴/Δ⁴^* embryos, we did not find evidence of global changes to the examined epigenetic marks deposited by these differentially expressed genes. These findings suggest that a robust network of epigenetic regulators may compensate for Ehmt2 loss of function and permit embryonic development and survival in *ehmt2^Δ⁴/Δ⁴^* mutant zebrafish.

## DISCUSSION

Our findings represent, to our knowledge, the first reported germline zebrafish model for Ehmt2 loss of function. We found that *ehmt2^Δ⁴/Δ⁴^* embryos displayed reduced *ehmt2* gene expression and global reduction of H3K9me2 marks. Additionally, we found that discrete H3K9me2 peaks normally present in promoter regions at 72hpf are absent in *ehmt2* mutants. When we examined how gene expression is altered under Ehmt2 loss of function, we identified two expression profiles of interest. The first cluster of genes were of comparable expression levels at the earlier timepoint (24hpf) in mutant and control embryos, however, at 48hpf, these genes would normally be downregulated in control embryos but instead showed sustained expression in mutant embryos. Genes within this cluster possessed key functions in developmental processes, particularly in neurodevelopment, as well as in several regulatory processes. The second cluster identified had genes which were downregulated in mutants as early as 24hpf compared to control, and the expression differences between mutant and control embryos grew by 48hpf for these genes. Together, these two expression profiles provided evidence for a putative growth defect in *ehmt2* mutants.

By tracking embryos through early development, we found mutant embryos fell behind as early as the 128-cell stage, but the variation in timing between mutant and WT embryos became most pronounced around the sphere stage of development. Given the sustained upregulation of neurodevelopmental genes in our RNA-Seq data, and the known functions of Ehmt2 in neural tissues, we examined mutant retinas and found that, even in staged-matched embryos, the mutant retina was smaller than the control retina at early timepoints. Interestingly, we found the size discrepancy between mutant and control retinas was negligible by around 72hpf. Consistent with these findings, when we probed cell cycle dynamics, we found a delayed peak in proliferation in the retina. This delayed proliferation was coupled with slower cell cycle progression between S-M. Finally, we probed how mutants may compensate for Ehmt2 loss of function by examining other chromatin modifiers. Of the modifiers which showed significant expression changes between mutants and control, we found the majority of chromatin remodellers and epigenetic modifiers associated with active transcription showed sustained upregulation in mutants. Despite significant changes in expression of several modifiers, we did not find global changes to two of the histone marks these genes were known to deposit in mutants (H3K27me3, H3K4me3), though this does not prove gene/region-specific changes are absent.

Ehmt2 appears to have conserved and varied roles in a variety of tissues across vertebrate species. In addition to its canonical role in development by mediating gene silencing *via* H3K9me2, several other roles are known to be affected in current Ehmt2 loss of function models. Our gene expression analyses have shown consistency with published findings. For instance, aberrant expression of DNA replication and repair machinery has been reported in Ehmt2 loss of function, where researchers showed Ehmt2 stabilizes the replication fork, maintains chromatin on newly synthesized DNA, and suppresses endogenous DNA damage [7,13,22]. Furthermore, many differentially expressed genes in our data are involved in neural pathways (neural regulators, neurotransmission, cell migration etc), and Ehmt2 is known to possess a multifaceted role in neural tissues. We showed retinal developmental delay was a function of slower cell cycle progression of retinal progenitor cells. The misregulation of DNA replication and repair genes, altered genome topography due to global H3K9me2 loss, and sustained expression of neural progenitor genes *via* loss of gene-specific H3K9me2 peaks may all promote prolonged cell cycle phases. Further research into distinct mechanisms will be required to tease apart whether Ehmt2 function is more crucial for neural development than the others, particularly through examining how global H3K9me2 loss affects chromatin accessibility and integrity. Additionally, given the role of EHMT in regulating several behavioural processes in *Drosophila*, and the neural developmental delay in *ehmt2^Δ⁴/Δ⁴^* mutants, it would be interesting to examine how germline Ehmt2 loss of function in zebrafish may affect neural and behavioural processes.

Ehmt2 loss of function in mice is embryonic lethal. This lethality has been explained by global H3K9me2 loss causing over-activation of cis-regulatory and repetitive elements resulting in ineffective regulation of early differentiation programs [36]. In contrast, *EHMT* loss of function mutant *Drosophila* are assumed to be adult viable and fertile since they do not show global H3K9me2 reduction [16–18]. *C. elegans* models which are deficient for H3K9 methyltransferases challenge this hypothesis, since they produce (temperature-dependent) viable embryos despite global loss of H3K9me2/3. However, the adult worms were sterile due to DNA-damage-related apoptosis from increased mutations and replication stress [19,24]. We were intrigued to find that *ehmt2^Δ⁴/Δ⁴^* fish were adult viable and fertile despite a global reduction in H3K9me2. The embryonic delay we observed in *ehmt2* mutant embryos was apparent by the sphere stage, at which point zebrafish embryos are normally undergoing the process of zygotic genome activation [37]. Given that H3K9me3 enrichment increases massively during this process [38,39], it may be that Ehmt2-mediated H3K9me2 loss prevents efficient H3K9me3 deposition and/or zygotic genome activation in our mutants, which could explain the appearance of delayed development around this timepoint. The rapid onset of developmental variation in *ehmt2^Δ⁴/Δ⁴^* embryos that does not appear to worsen progressively over time further supports this hypothesis, rather than the alternative explanation that changes to differentiation programs (as predicted in mice) plays the predominant role in developmental delay. It should be noted that the developmental delay we have shown, while significant, is relatively mild. We might expect the fitness of these embryos to be called into question under challenging circumstances outside of highly controlled laboratory conditions, especially considering the intra-clutch variation observed. It would be interesting for future work to examine how these embryos handle environmental pressure such as changes to temperature/salinity, and whether they can still meet all developmental milestones.

In this study, we demonstrated a new Ehmt2 germline loss of function model using zebrafish. These mutants exhibit a mosaic molecular phenotype of known Ehmt2 loss of function models, with embryonic survivability despite a global reduction in H3K9me2. Our findings suggest that the lack of normal H3K9me2 deposition in early timepoints results in embryo-wide developmental delay, while the additional retinal delay (witnessed in stage-matched embryos) is a consequence of Ehmt2-dependent neural gene regulation. The mechanisms enabling *ehmt2^Δ⁴/Δ⁴^* embryos to progress through the cell cycle and ultimately survive to adulthood, remains to be determined. While we found gene expression changes of various histone modifiers, for the canonical functions of these modifiers that we probed, we found no significant global changes to their histone marks. This may be due to genomic redundancy, or altered recruitment pathways promoting the robustness in epigenetic regulation [40]. Regardless, our novel mutant zebrafish model provides a promising avenue for examining the global impacts of Ehmt2 loss of function on the genomic regulation of major developmental processes.

## MATERIALS AND METHODS

All materials and reagent information is described in S1 Table.

### Ethics Statement

All animals were treated in accordance with the regulations on animal experimentation established by the Canadian Council on Animal Care (CCAC). The experimental procedures were approved by the University of Toronto Animal Care Committee.

### Zebrafish Husbandry & embryo culture

All zebrafish strains were used in this study were housed in a recirculation system with 14hr–10hr light-dark cycles at a constant 28 °C. The *Tg(h2af/z:GFP)* strain [41] was a kind gift from Dr. Ian Scott (Hospital for Sick Children, Toronto), and the *Tg[EF1a:mAG-zGem(1/100)^rw0410h]* strain [34] was provided by the laboratory of Dr. Atsushi Miyawaki from the Zebrafish National BioResource Project. All mutant fish used in this study were derived from the wildtype-AB zebrafish strain background obtained from the Zebrafish International Resource Center (ZIRC) and maintained at our facility. Two *ehmt2* mutant lines were generated: the 4bp deletion mutant described in this study, and a mutant with a 69bp deletion in the same region (data not shown). Characterisation of both mutants showed the same phenotype. Since both *ehmt2^Δ⁴^* and *ehmt2^Δ⁶⁹^* homozygous mutants survived to adulthood, they could be maintained as adults for breeding. They have been maintained in our lab for >5 generations and embryos have consistently displayed the same phenotype over multiple generations.

Unless otherwise stated, embryos were collected and grown in a (dark) 28 °C incubator, in a 100 mm wide petri dish in facility water, until the required developmental stage (up to 3 days post fertilisation). Sexual dimorphism is not detectable in zebrafish before roughly 2–3 months post fertilization, and fish do not have specific sex chromosomes that can be used for genotyping. Therefore, sex as a biological variable was not considered in this study.

### Embryo Manipulations

Injections of antisense morpholino constructs was performed at the 1 cell stage into the yolk adjacent to the embryo. Successful incorporation of the construct was judged by Fast Green staining incorporated into the injection mix. The Ehmt2 inhibitor Unc0638 was purchased from Millipore Sigma (U4885), and the A-366 compound [42] was provided by the Structural Genomics Consortium, Toronto. Following dose-response curves (data not shown), embryos were treated with 50µM Unc0638, 25µM A-366, or DMSO as a carrier control. Embryos were treated on day 0 after removal of the chorion (approximately 2-3hpf) and cultured overnight in the dark at 28°C in facility water and inhibitor. Embryos were washed with facility water and treated with fresh inhibitor at 24-hour increments until fixation.

### Generation of CRISPR-Cas9 mutants

The protocol used in this study for CRISPR-Cas9 genome editing was previously described in Olsen *et al.* [43]. Briefly, equal concentrations of sgRNA and Cas9 mRNA (400-600ng, transcribed from pT3TS-nCas9n plasmid (Addgene #46757, RRID: Addgene_46757) were co-injected into the cell (through the yolk) of a 1-cell staged embryo and grown to adulthood. Several Guide RNAs (and PCR primers) were designed to target exon 5 of *ehmt2* using CHOPCHOP [44]. The final target sequence, along with the PCR primers and restriction enzyme used for sequencing were as follows:

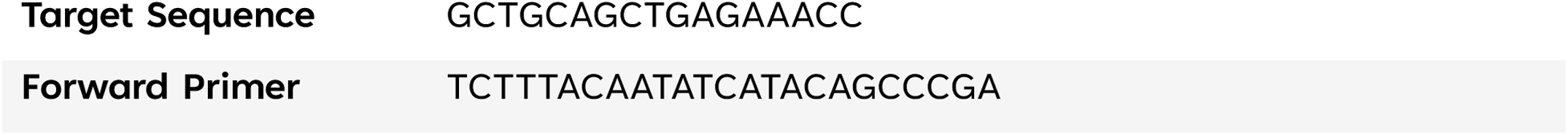
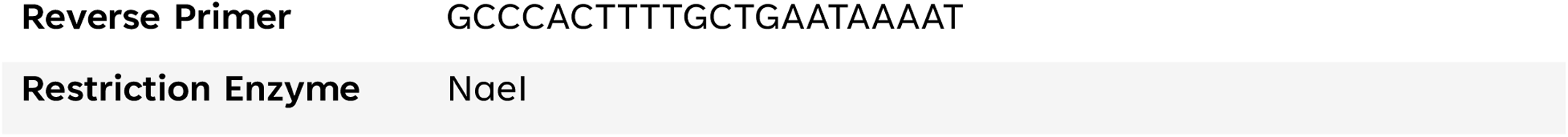

The mutant lines identified and used for further analyses had a 4bp deletion (*ehmt2^Δ⁴^*) and a 69bp deletion (*ehmt2^Δ⁶⁹^*, data not shown) in the same site. All mutations were examined by fin clipping anesthetised (tricaine), adult fish, followed by PCR, RE digest (as RE site was eliminated in mutants) and gel electrophoresis. The mutations were also validated through sequencing of the PCR product. The *ehmt2 ^Δ⁴^* line was maintained as adult homozygous mutant (*ehmt2^Δ⁴/Δ⁴^*) and WT siblings (*ehmt2^+/+^*) (progeny of heterozygous in-cross). For experimentation, homozygous mutants were in-crossed and the resulting embryos used as ‘mutant’ embryos, while the progeny of WT sibling in-crosses were used as ‘control’ embryos.

### Embryo Staging

Live embryos were staged according to the developmental stage descriptions outlined by Kimmel *et al.*[45]. Briefly, 4-cell staged embryos were placed into individual wells of a 24-well plate in 1 mL of facility water and the time recorded. The plate was kept in a 28 °C incubator (in the dark), and embryos were periodically checked for developmental stage using an upright light microscope. The stages selected to record were 128-cell, sphere, 30% epiboly, shield, 80% epiboly, 22-somite, and prim-5. These were selected based on the ability to differentiate that stage from those around them. Ten control and ten mutant embryos were tested per replicate.

### Immunofluorescent Staining

Embryos were fixed in paraformaldehyde (4% w/v, 4 overnight), cryo-sectioned and processed for staining as previously described [43,46]. An antigen retrieval reaction was performed on sections being stained PCNA and PHH3 by incubating sections in Sodium Citrate (10mM, 0.05% Tween, pH 6.0) at 95°C for 30min, cooling and washing 3 times in PBST (0.2% Tween in PBS). Antigen retrieval reactions were not required for any other antibody staining. Samples were blocked in PBST with Normal Goat Serum (2%) for primary antibody overnight at 4°C in a humid chamber. Once primary staining was completed, tissues were washed in PBST, counterstained with Hoechst 33342, rinsed thoroughly in PBS (1X), cover slipped with 90% glycerol in PBS, sealed with clear nail polish and stored at 4°C until imaging. Antibodies and conditions used are listed in Table S1.

### Acridine Orange Staining

Acridine orange was used as a rapid method for detecting cell viability. Embryos were stained just prior to mounting and imaging. Embryos were placed in Acridine orange solution (1µg/ml Acridine orange in facility water) for 1 hour, then washed 3 x 5 min in facility water before being immobilised on their side in 1% agarose in a glass bottom dish. During imaging, z-stack images were generated throughout the entire retina (from 80 µm slices) and the number of acridine orange-positive cells counted. n = 12 (4 embryos x 3 separate clutches).

### Live Imaging of Transgenic Embryos

*Tg(h2af/z:GFP)* and *Tg[EF1a:mAG-zGem(1/100)^rw0410h]* fish were independently outcrossed with mutants and bred to homozygosity. Whole mount live confocal imaging of embryos was accomplished by anaesthetizing embryos in tricaine (0.16 g/L) approximately 30 minutes before imaging, and immobilising them on their side in 1% agarose in a glass bottom dish. Tg[h2af/z:GFP]-positive embryos were imaged for three hours between 21 and 24 hpf at 5 min intervals. Mitosis time was calculated by measuring the time from nuclear condensation to separation into two distinct nuclei*. Tg[EF1a:mAG-zGem(1/100)^rw0410h]*-positive embryos were imaged for seven hours between 20 and 27hpf at 5 min intervals. Interkinetic migration during G2 was measured from the time a nucleus began moving from the basal side of the lamina to when it stopped moving near the apical side. The apical dwell time was then measured until condensation of nucleus occurred, marking the entry into mitosis.

### Cell Cycle Analysis by Flow Cytometry

Sample preparation for Flow Cytometry analysis was adapted from Quillien *et al.* [47]. Single cells were isolated from 24hpf mutant and control embryos. Pools containing 25 embryos were disaggregated in TryplE (1.3ml), and the resultant solution was passed through a 70µm mesh into PBS (1X, 3ml). Biotium Live-or-Dye 640/666 (4µl) was added and the cells incubated in the dark at 4°C for 30min with inversion every 5 min. Cells were spun down at 300g for 5min and the supernatant removed. Cells were then fixed in cold PFA (4% in PBS, 1ml) at 4°C for 30min. Cells were spun down at 3,000g for 5min and the supernatant removed. Cells were washed in cold PBS (1X, 1ml), spun down, and the supernatant removed. Each sample was resuspended in 250µl of sort buffer (1ug/ml DAPI in BSA (1% in 1X PBS)) and strained through a 35µm mesh. Samples were stored on ice until analysis. Flow Cytometry was performed on a BD LSRFortessa™ X-20 and cell stage was determined *via* chromatin content determined by DAPI incorporation [48]. Data was analysed using OMIQ software (www.omiq.ai) from Dotmatics (www.dotmatics.com).

### ChIP Sequencing

The protocol used here was modified from Kron *et al.* 2017 [49]. Beads were prepared combining equal parts of Protein A and Protein G Dyna beads, followed by washing thrice with PBS-BSA (5 mg/ml BSA). H3K9me2 antibody (3 µg per sample) in PBS-BSA were added to the washed beads and agitated overnight at 4°C. Mutant and control samples were collected in triplicate (3 individual clutches) at 24 and 72 hpf. For each sample, 50 embryos were collected from a single clutch and fixed in PFA (4% w/v) for 2 hours at room temperature, followed by washing with PBS (1X). Samples were crosslinked with formaldehyde (1% w/v), followed by addition of Glycine (to a final concentration of 0.125M) for 5 min to halt crosslinking. Samples were then washed with cold protease inhibitor in PBS, then cold protease inhibitor in PBS-BSA. Embryos were homogenised with a pestle in lysis buffer (1% w/v SDS, 10mM EDTA, 50mM Tris HCl pH8, 1X protease inhibitor) then sonicated for 28 cycles (30 sec on, 30 sec off). Following centrifugation, 30 µl of supernatant from each sample were retained and stored as input samples. Dilution buffer (1% v/v Triton, 2mM EDTA, 20mM Tris HCl pH8, 150mM NaCl, 1X protease inhibitor) and beads were added to remaining supernatant, and samples agitated overnight at 4°C. Samples were next aspirated and treated with the series of buffers described below:

|  |  |
| --- | --- |
| **1) low salt buffer** | 0.1% w/v SDS, 10mM EDTA, 50mM Tris HCl pH8, 1% v/v Triton, 150mM NaCl |
| **2) high salt buffer** | 0.1% w/v SDS, 10mM EDTA, 50mM Tris HCl pH8, 1% v/v Triton, 500mM NaCl |
| **3) LiCl buffer** | 10mM Tris HCl pH8, 1mM EDTA pH8, 250mM LiCl, 0.5% v/v NP40, 0.5% w/v sodium deoxycholate |
| **4) TE buffer x 2** | 10 mM Tris-HCl, pH 8.0, 10 mM EDTA |

To both ChIP and input samples, de-crosslinking buffer (1% SDS, 0,1M NaHCO_3_) was added and samples agitated overnight at 65°C. Finally, samples were purified with a MinElute PCR Purification Kit (*following manufacturer’s instructions*) and eluted into 30 μl of nuclease free water.

DNA quality and fragment size were checked using an Agilent TapeStation and showed an average fragment length of 200bp. Samples were submitted to The Centre for Applied Genomics at the Hospital for Sick Children Sequencing Facility (Toronto, ON) for library preparation and sequencing. Library preparation was completed using a NEBNext Ultra II DNA Library Prep Kit for Illumina. Sequencing was performed on an Illumina NovaSeq 6000 – S4 flowcell for paired end 100bp sequencing at ∼50M reads per sample. Sample file information can be found in Table S2.

Sequencing analysis Reads were mapped against the *Danio Rerio* genome GRCz11 (GCF_000002035.6, NCBI) using Bowtie2 (v2.5, RRID:SCR_016368) [50]. File sorting and filtering was performed using SAMTOOLS (v1.14, RRID:SCR_002105) [51] and sambamba (v1.0, RRID:SCR_024328) [52]. Peak calling was performed with MACS2 (v2.2, RRID:SCR_013291) (with the additional parameter: “--broad”) [53], using the readily available genome size for *D. Rerio* genome GRCz10 (1.37×10^9^) from Deeptools (v3.51, RRID:SCR_016366) [54].

Quality assessment was performed using ChIPQC (v1.44, www.bioconductor.org), using a custom annotation generated from the same genome (GRCz11, GCF_000002035.6). Briefly, the FriP scores averaged 15% in mutant *vs* 13% in control indicating slightly better enrichment in the mutant samples, and there was increased variability observed in the 24hpf mutant replicates (Figure S6A & B). Cross correlation analysis indicated all samples had consistent fragment size with an absence of phantom peaks (Figure S6C). Peak profiling showed higher signal intensities for mutant compared to control samples, likely due increased chromatin accessibility in mutants from global H3K9me2 loss (Figure S6D).

DiffBind (v3.18, RRID:SCR_012918) was used to perform differential enrichment analysis with R Project for Statistical Computing (v4.5, RRID:SCR_001905) [55,56]. The broadPeak file type was used from the macs2 output. Briefly, the peaks were imported and merged into a consensus set for each sample type and then counts computed using the following parameter: bUseSummarizeOverlaps = TRUE. A correlation map was also plotted showing discrete sample clustering (Figure S7). Next differential enrichment analysis was performed (using method = DBA_ALL_METHODS). Principle component analysis (PCA) plotted using only significantly enriched peaks (Figure 1E) and a volcano plot was generated showing significantly enriched peaks in pink (Figure 1F). Annotation and functional enrichment was completed used ChIPseeker (v1.44, RRID:SCR_021322) [57] and biomaRt (v2.64, RRID:SCR_019214) [58].

### RNA Sequencing

Mutant and control samples were collected at 24 and 48 hpf. For each sample type, 48 embryos were pooled from mixed clutches (due to breeding issues following COVID-19 shutdowns) and split into 4 replicate groups. Total RNA was extracted from each sample using a Zymo Research Quick-RNA MicroPrep Kit (following the kit protocol) and eluted into 10 μL of nuclease free water. Samples were submitted to The Centre for Applied Genomics at the Hospital for Sick Children Sequencing Facility (Toronto, ON) for quality control, library preparation and sequencing. Briefly, RNA quality was checked using an Agilent BioAnalyzer, confirming RNA integrity numbers > 9, and that at least 80% of RNA was 200bp or longer, for all samples. Library preparation was completed using the NEBNext Ultra II Directional RNA Library Prep Kit for Illumina with the NEBNext Poly(A) mRNA Magnetic Isolation Module). Sequencing was performed on an Illumina NovaSeq 6000SP flowcell for paired end 100bp sequencing at ∼50M reads per sample. Sample file information can be found in S3 Table.

Reads were mapped against the *D. Rerio* transcriptome (GRCz11, Ensembl) using Kallisto (v0.46.2, RRID:SCR_016582) (see Figure S8A & B) [59]. Raw read data (data not shown) and mapped counts (Figure S8C & D) were quality checked using MultiQC and mean sequence quality scores were above 30 in all samples. The mapped reads were imported into RStudio (v 2021.9.0.351, RRID:SCR_000432) using tximport (v1.20, RRID:SCR_016752) and annotated using biomaRt. EdgeR (v4.6.2, RRID:SCR_012802) [60] was used to calculate counts per million (cpm), and the samples were filtered to remove genes with no counts before normalisation using EdgeR (method: Trimmed Mean of M-values (TMM) (Figure S9). Samples were clustered using varying hclust and distance methods to confirm discrete clustering of different sample types (Figure S10A). Principle component analysis was performed and visualised using R package: stats-package (v4.5, RRID:SCR_025678). Heteroscedasticity of RNA-Seq data results in increase variance with increased mean, so LIMMA (v3.64.1, RRID:SCR_010943) [61] was used to variant-stabilise (voom [62]) the data and calculate Bayesian statistics (eBayes) (see Figure S10B). Volcano plots were generated using ggplot2 (v3.5.2, RRID:SCR_014601) [63]. Differentially expressed genes were defined as those with a fold change of ±2 (between genotypes) and an adjusted p-value of less than 0.05. Unsupervised clustering was performed using Clust (v1.12, RRID:SCR_025283) (using the parameter “-n 101 4”) [20]. Functional enrichment was performed using gProfiler2 (v0.2.3, RRID:SCR_018190) [64] on the top 500 ranked genes from a linear model fit (limma, topTable with method = “BH”). Unless otherwise stated, plots and heatmaps were generated using ggplot2. RNA-Seq results were validated by sampling several differentially expressed genes using RT-qPCR using *actin* expression for normalisation (Figure S11).

### RT-qPCR

Pools of 20 embryos from a single clutch were collected for each sample replicate. Total RNA was extracted from each sample using a Zymo Research Quick-RNA MicroPrep Kit (following the kit protocol) and eluted into 15 μL of nuclease free water. RNA integrity and quantification was examined using a NanoDrop and equal concentrations used for each sample moving forward. RNA was reverse transcribed using SuperScript IV Reverse Transcriptase according to the manufacturer’s instructions. Primers were designed using PrimerQuest (Integrated DNA Technologies, USA, www.idtdna.com). RT-qPCR was completed on three biological replicates (in technical triplicates) using SsoAdvanced Universal SYBR Green Supermix and the following primers:

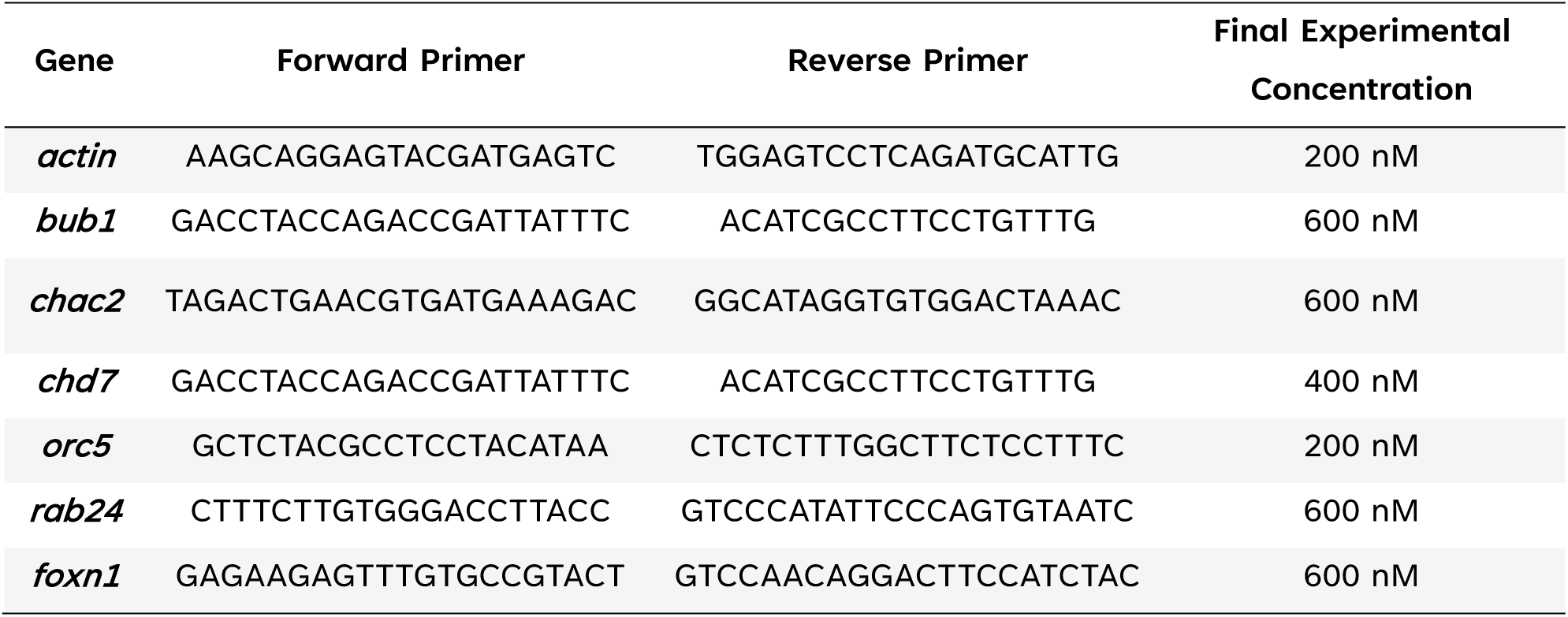

The qPCR was run on a CFX OPUS 384 Real-Time PCR system running CFX Maestro (v2.3, Bio-Rad Laboratories, Inc., USA) using the following conditions: 1) 95°C for 5 min, for 40 cycles: 95°C for 10 sec, 60°C for 30 sec 3) melt curve 65 to 95°C ramped at 0.5°C/5 sec.

### Imaging & Data Processing

Unless where otherwise specified, imaging was completed using Leica TCS SP8 confocal microscope. Images were processed using Leica Application Suite (LAS X) (v3.7.6, RID:SCR_01367) and FIJI[65] software programs. Photographs of adult zebrafish were captured using a LUMIX G9 camera. Fluorescence quantification and cell counting were completed using Imaris (v10.2, RRID:SCR_007370), by averaging relative fluorescence or cell counts, respectively, from three sections (20 µm) of the central retina. Except for where described in sequencing analyses, statistical analyses were completed using RStudio[55,56] and/or Microsoft Excel (RRID:SCR_016137). Unless otherwise stated, statistical significance was calculated using a one-way ANOVA.

## ACKNOWLEDGEMENTS

We thank Karen Ho, Dr. Valerie Crowley, and the team at The Centre for Applied Genomics at the Hospital for Sick Children Sequencing Facility (Toronto, ON) for assistance with library preparation and sequencing of RNA-Seq and ChIP-Seq samples. We thank Dr. Giacomo Grillo for advice on ChIP-Seq sample preparation. We thank Dr Nathalie Simard and Vincent Cheng from the Temerty Faculty of Medicine Flow Cytometry Facility at the University of Toronto (Toronto, ON) for advice and assistance in gathering flow cytometry data. We thank Dr. Dalia Barsyte-Lovejoy from the Structural Genomics Consortium for sharing reagents. This research was funded by a Discovery Grant from NSERC and the Department of Cell & Systems Biology, University of Toronto (V.T.).

## DATA AND MATERIALS AVAILABILITY

Processed data are available in the main text or the supplementary materials, raw sequencing data will be made publicly available upon journal submission.

