## Supplemental Figures for "Loss of Ehmt2/G9a function in zebrafish is associated with global deficiency in H3K9 dimethylation, misregulated cell cycle dynamics, and embryonic developmental delay"

### SUPPORTING INFORMATION

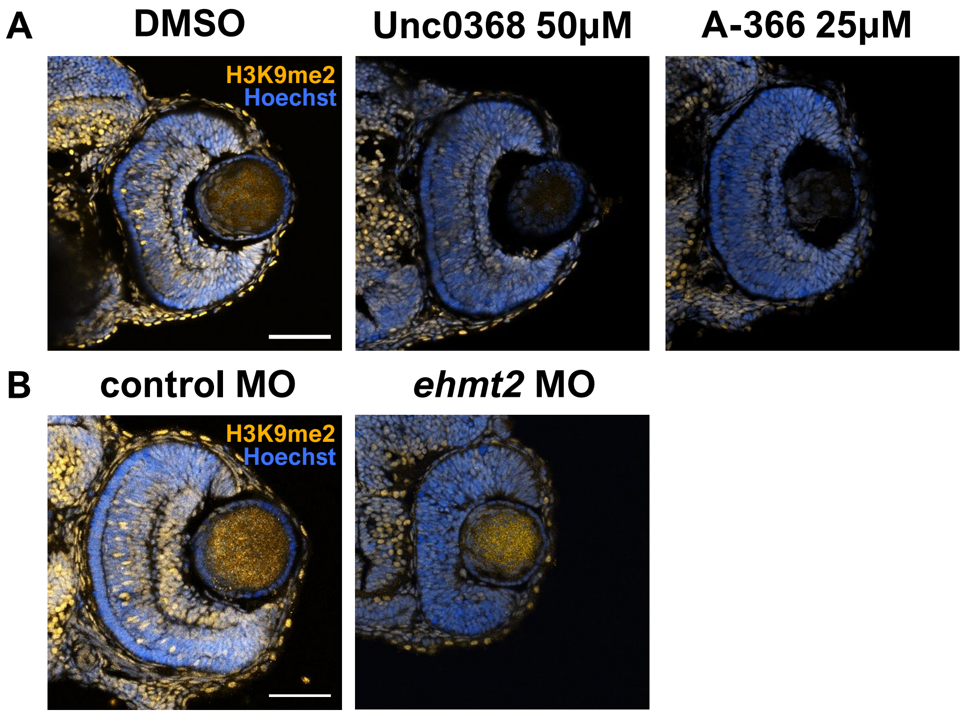


**Figure S1: Ehmt2 knock down embryos show global reduction of H3K9me2.** Immunostaining against H3K9me2 on retinal sections for drug-treated embryos (A) and morphants (B) at 48hpf. Scale bars = 50 µm.

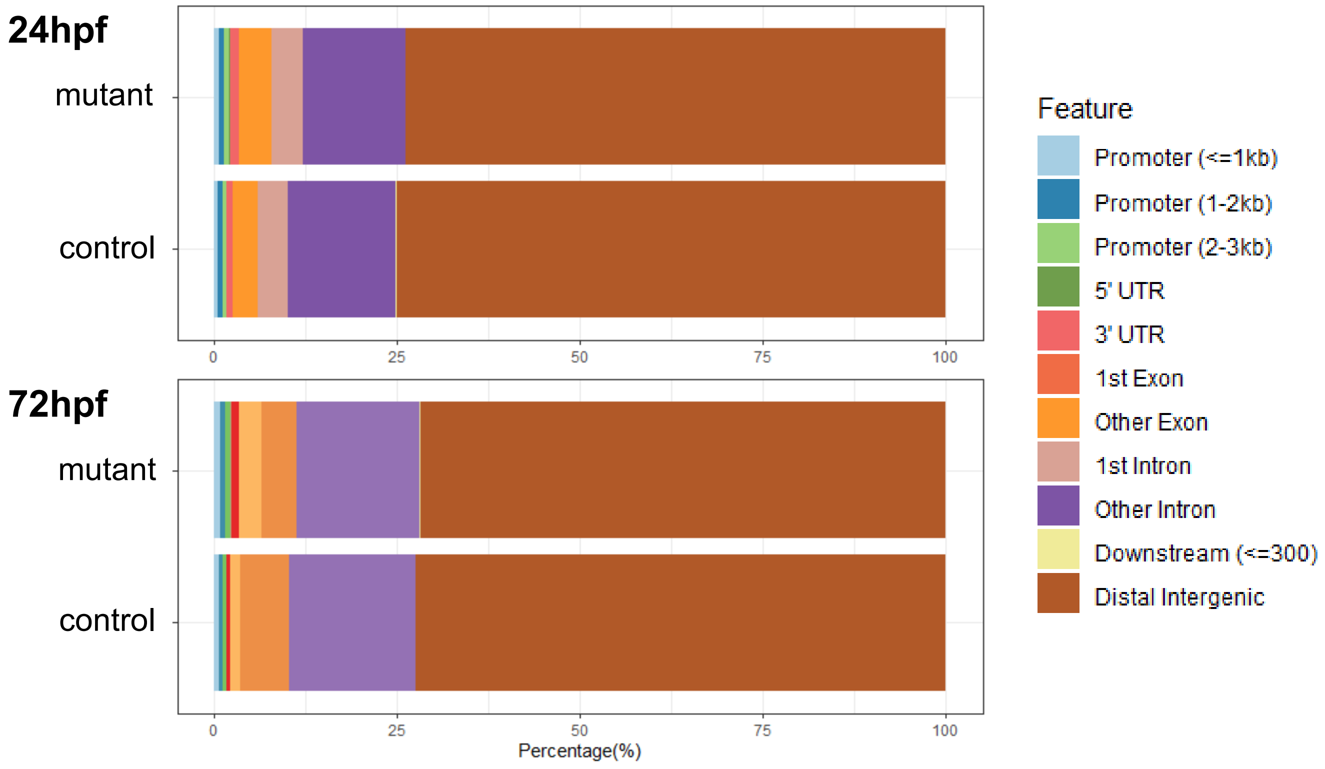


**Figure S2: Chip-seq results corroborate previous studies indicating the majority of H3K9me2 peaks occur in distal intergenic regions.** Plot showing distribution of H3K9me2 peaks across different genomic features.

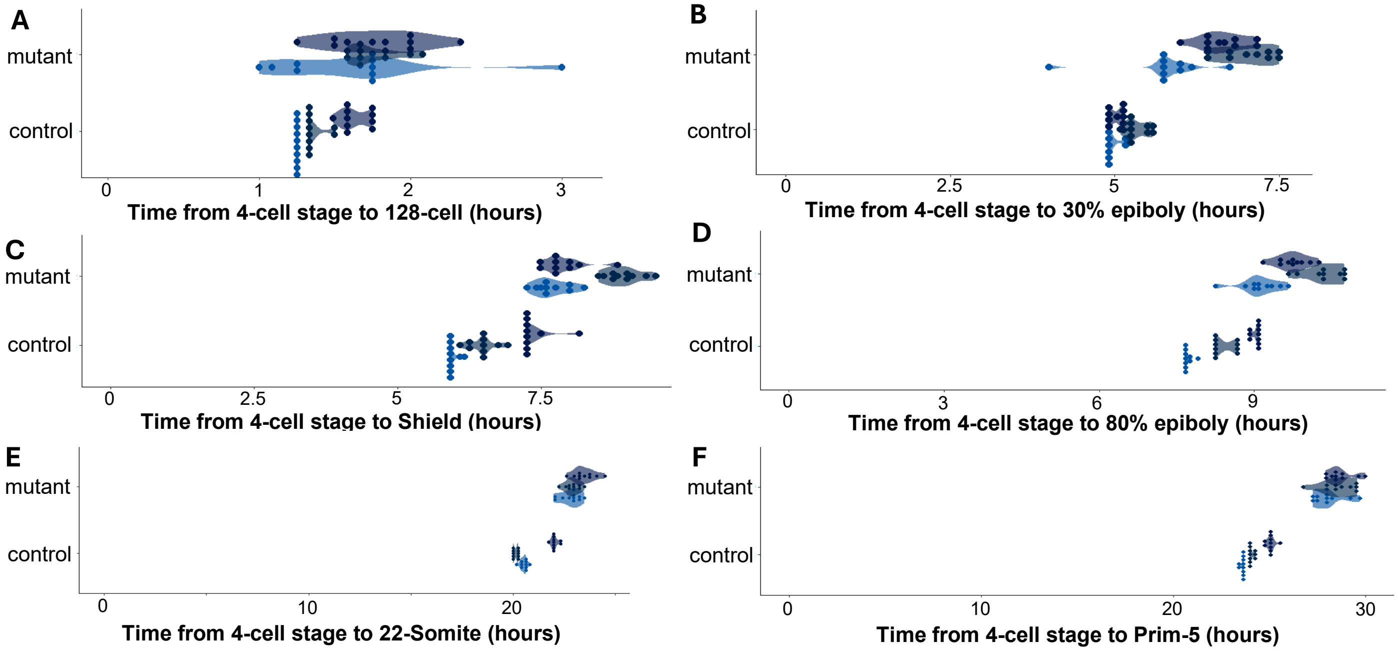


**Figure S3: *Ehmt2* mutants show increased intra-clutch variation.** Individual stage plots from Figure 3A sorted by replicate (n = 10), showing developmental time to 128-cell (A), 30% epiboly (B), shield (C), 80% epiboly (D), 22-somite (E), and prim-5 (F).

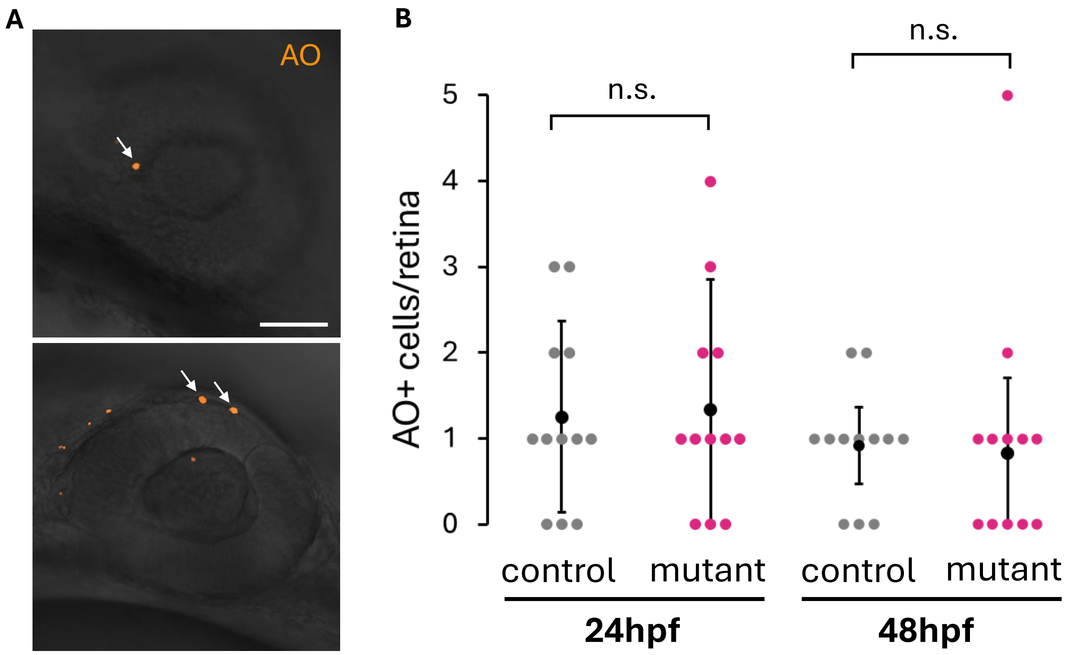


**Figure S4: *Ehmt2* mutants do not show increased cell death**. A) Example acridine orange (AO) staining in control retinas at 24 hpf (arrows indicate acridine orange-positive retinal cells, scale bar = 50 μm). B) Comparison of the number of acridine orange-positive cells per retina in mutant and control retinas at 24 and 48 hpf, n = 12.

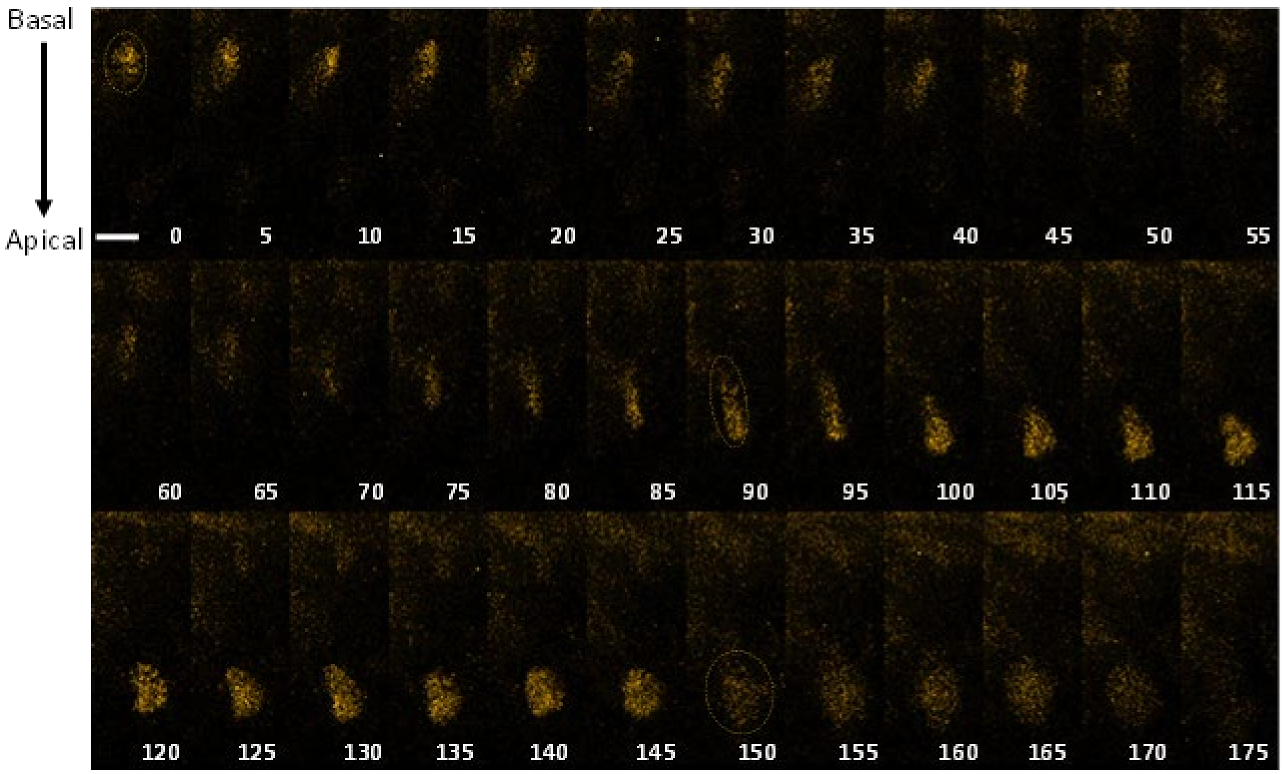


**Figure S5: Full time series of a single mAG+ retinal progenitor nuclei undergoing interkinetic nuclear migration in concordance with G2.** Time in min, scale bar = 10 µm.

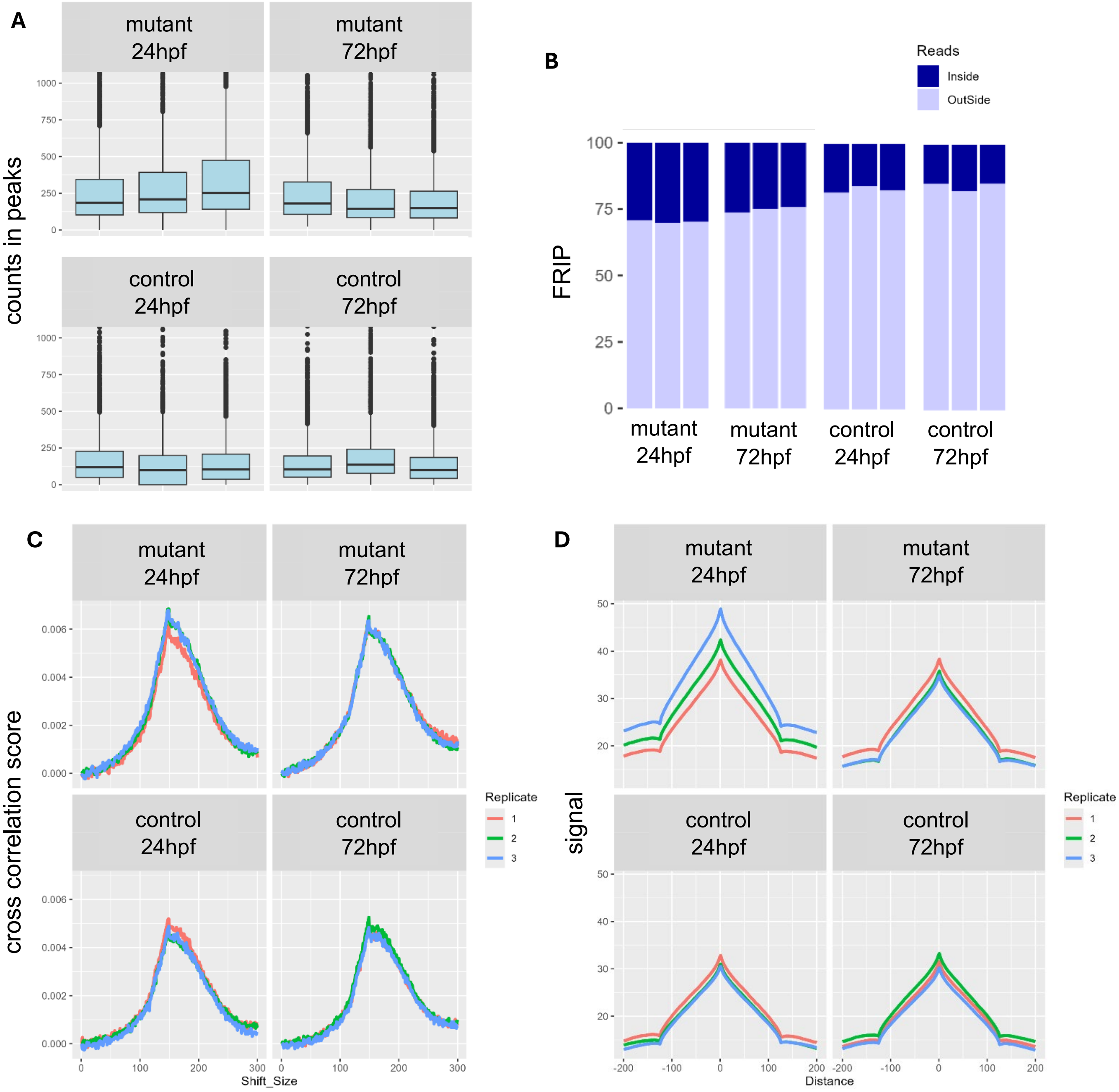


**Figure S6: ChIP-Seq quality control indicated good sample quality, with increased sequencing depth in mutants at both 24 and 72 hpf.** Read counts in peaks (A) and the fraction of reads in peaks (B) are similar between replicates. C) Plots of cross correlation analysis showed a single peak for each sample indicating consistent read lengths. D) Higher signal intensity was found in from the peak profiling indicate increased sequencing depth in mutant samples. All samples plotted by replicate.

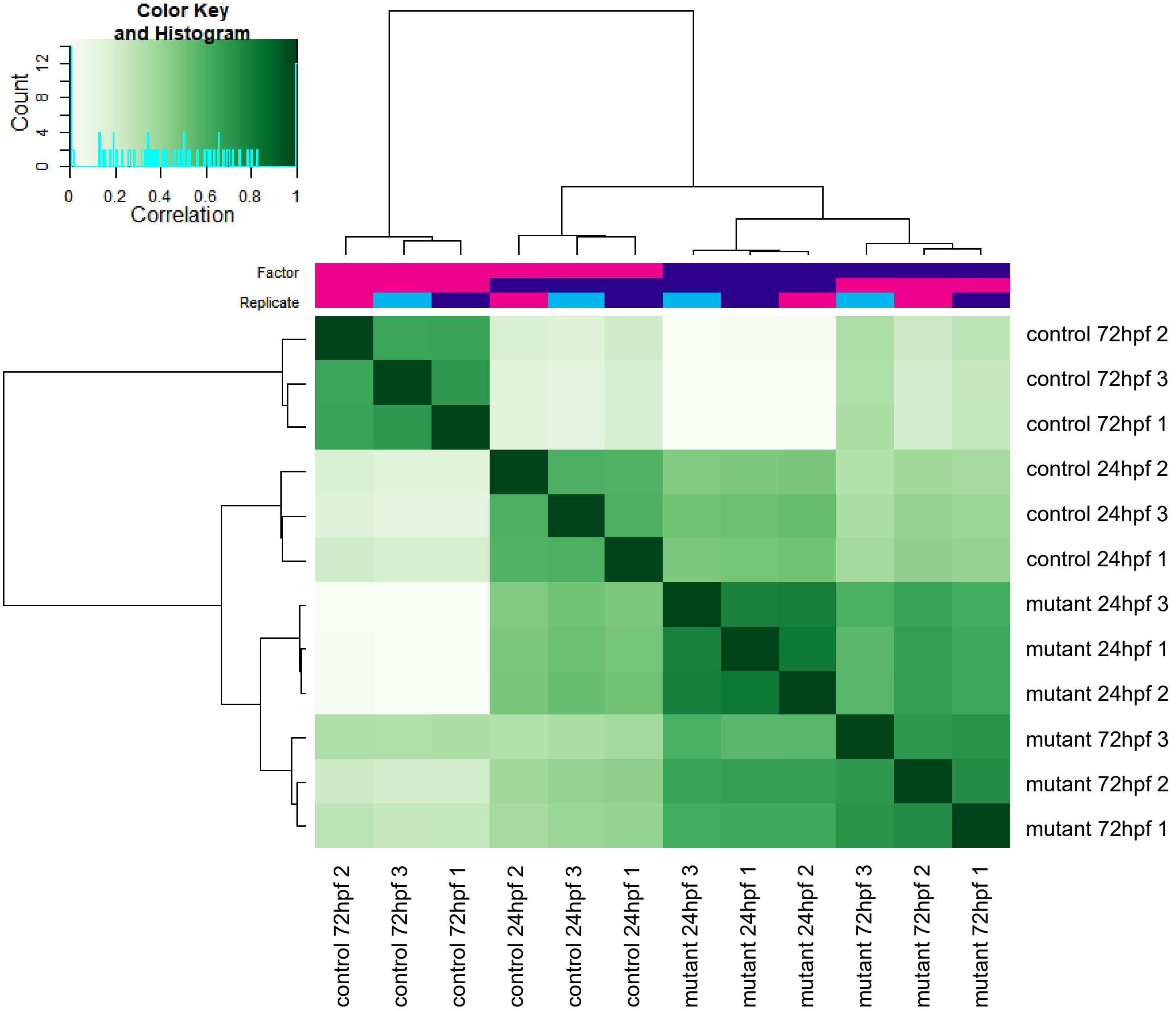


**Figure S7: Correlation heatmap of ChIP-Seq samples shows discrete clustering of samples types.**

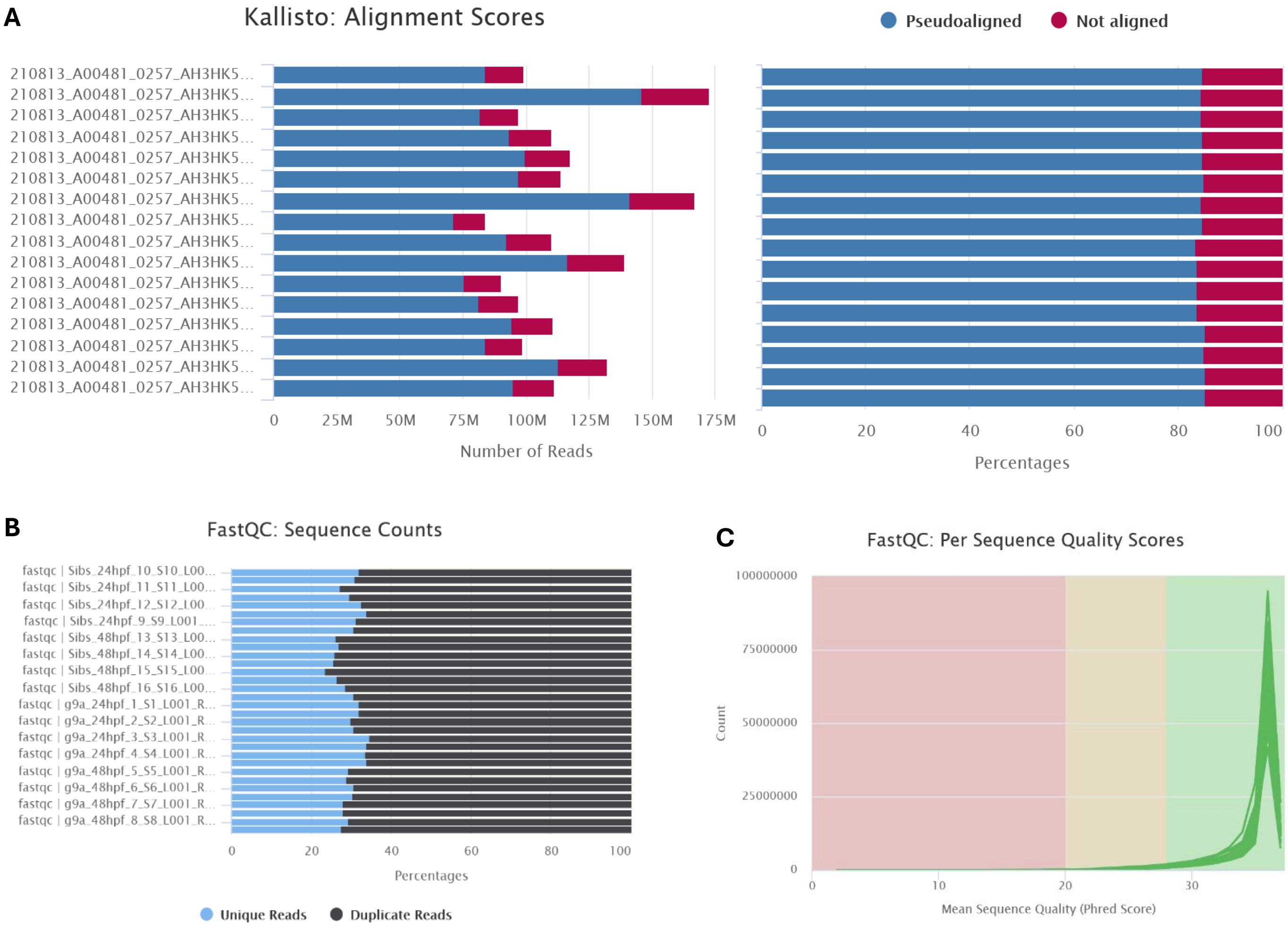


**Figure S8: RNA-Seq alignment and quality scores.** A) Kallisto alignment scores by number of reads (left) and as percentage of reads (right) showing similar patterns of mapped *vs* unmapped reads. B) FastQC sequence counts showing similar numbers of unique *vs* duplicated reads between samples. C) Per sequence quality scores are consistent for all samples.

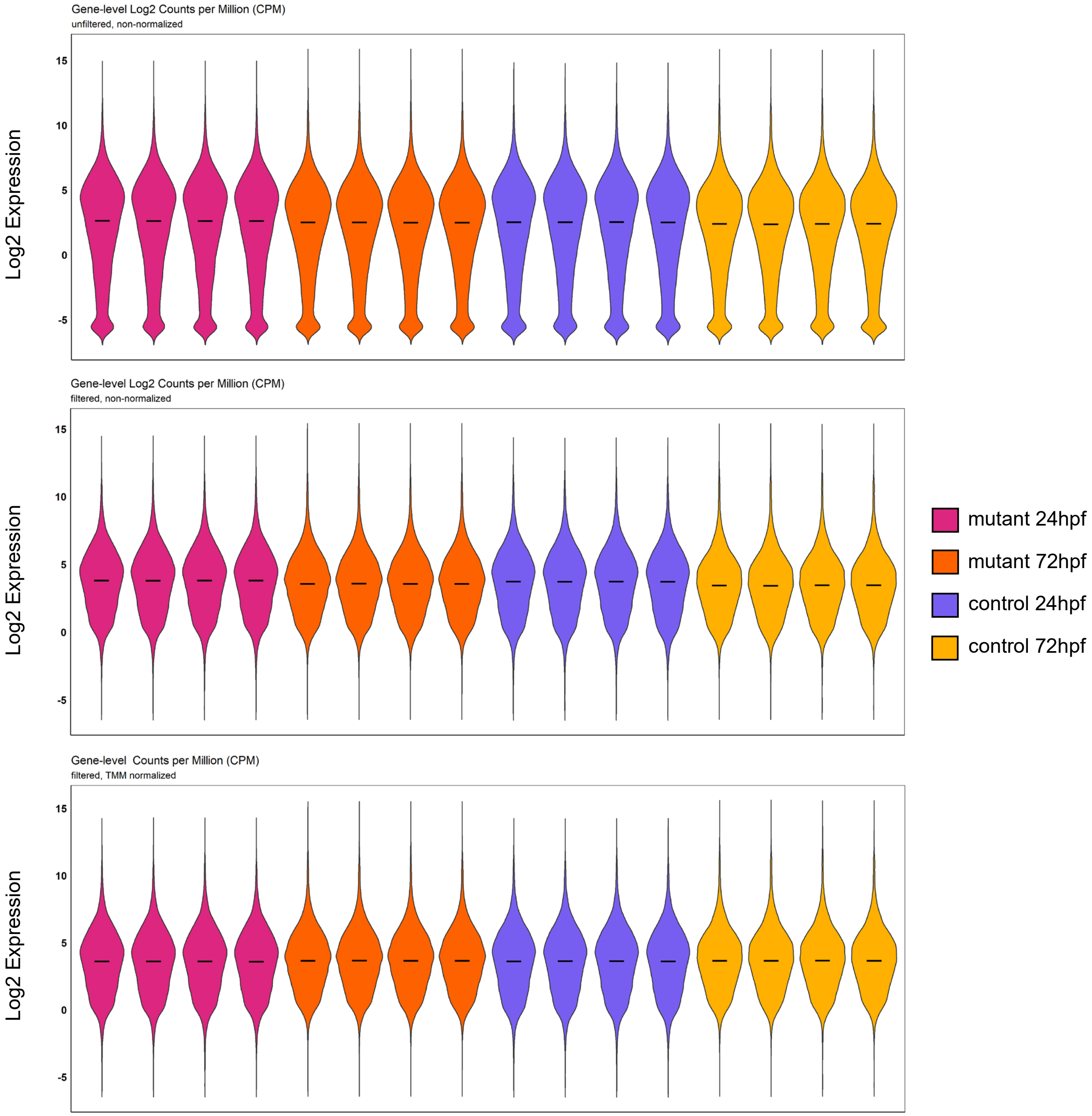


**Figure S9: Filtering and normalisation of RNA-Seq Data.** Plots of sample spread (as Log2(CPM)) prior to filtering (top), after filtering to remove zero-counts (middle), and after TMM normalisation. CPM = counts per million, TMM = trimmed mean of M-values. All samples plotted by replicate.

**A**

**B**

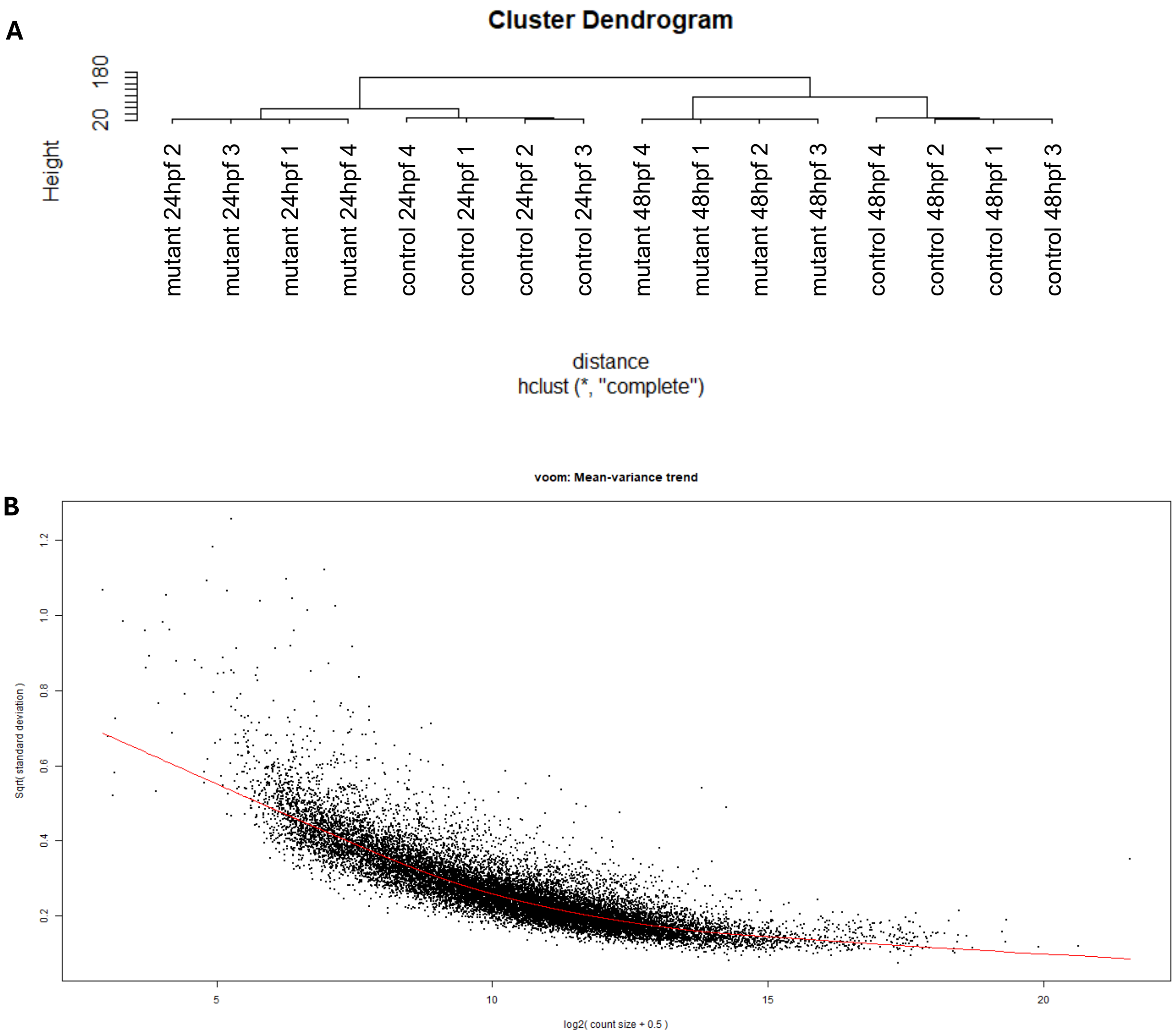


**S10 Figure: Clustering of RNA-Seq Samples and modelling of mean variance.** A) Example cluster dendrogram of RNA-Seq samples (generated using distance ='euclidean' and hclust method = 'complete'). B) Plot of mean-variance trend following variant-stabilisation with voom.

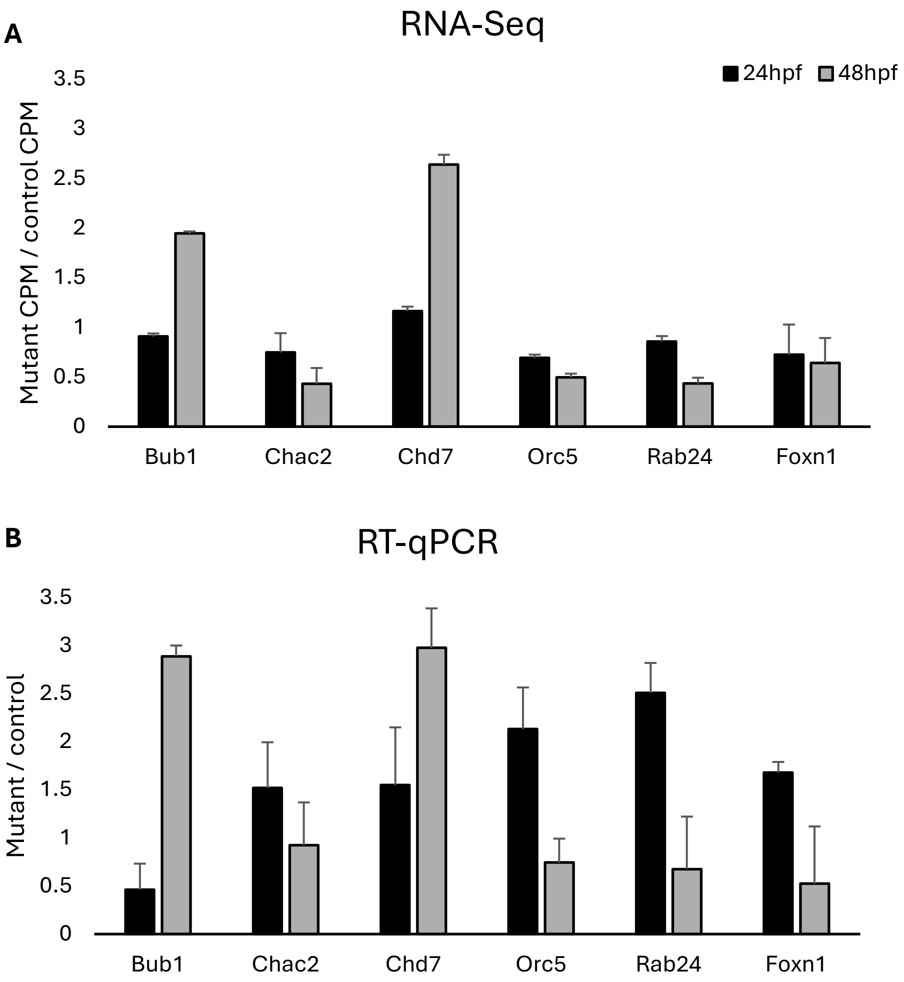


**Figure S11: RNA-Seq Validation.** Expression of six unrelated genes which exhibited differential expression between mutant and control embryos from 24 - 48 hpf in our RNA-Seq dataset (A) were tested via RT-qPCR (B) to show the same trends in gene expression. RT-qPCR scores were first normalised to *actin*. RNA-Seq n = 3, RT-qPCR n = 4, error bars = 1 standard deviation.
